# 10,239 whole genomes with multiomic and clinical health information as the Korean Multiomics Reference dataset

**DOI:** 10.1101/2025.11.17.688763

**Authors:** Kyungwhan An, Sungwon Jeon, Yoonsung Kwon, Yookyung Choi, Changhan Yoon, Yeonsu Jeon, Jihun Bhak, Dong-Hyun Shin, Hyoung-Jin Choi, Hyomin Lee, Yeo Jin Kim, Eun-Seok Shin, Hyojung Ryu, Asta Blazyte, Sangsoo Park, Juok Cho, Dan Bolser, Soobok Joe, Jin Ok Yang, Jongbum Jeon, Jong-Hwan Kim, Jungeun Kim, Dooyoung Jung, Yun Sung Cho, Kiyuk Chang, Eun Ho Choo, Eunmin Kim, Sang Yeub Lee, Weon Kim, Min Gyu Kang, Ae-Young Her, Suk Chon, Jeong-Taek Woo, Sang Youl Rhee, Siwoo Lee, Hyo-Jeong Ban, Hee-Jeong Jin, Younghwa Baek, Yong Min Ahn, Sang Jin Rhee, Min Ji Kim, Sang Yeol Lee, Chan-Mo Yang, Se-Hoon Shim, Seong-Jin Cho, Shin Gyeom Kim, Hyung-Tae Jung, Byung-Joo Ham, Yoon Young Choi, Jae-Ho Cheong, Seung-Ki Kim, Ji Hoon Phi, Seung Ah Choi, Heon Yung Gee, Sun Young Joo, Jinsei Jung, Wonsuk Shin, Sang-Hyuk Lee, Borah Kim, Woojae Myung, Chong Kun Cheon, Dong Uk Kim, Seok-Soo Byun, Gangnam Jin, Hojune Lee, Kyun Shik Chae, Chang Geun Kim, Bonghee Lee, Jaesuk Lee, Kwangwoo Kim, Semin Lee, Neung-Hwa Park, Haeyoung Jeong, George M. Church, Jong Bhak

## Abstract

We present Korea10K, the largest genomic dataset of the Korean population, comprising 10,239 high-coverage whole genomes (mean depth 30×) with matched multiomic profiles and phenotype data. Korea10K achieves complete and near-complete discovery of very rare and ultra-rare alleles, respectively, at 9,000 Korean genomes. This dataset provides the high-quality population-specific imputation panel, enabling accurate inference of low-frequency variants. Admixture analyses confirm the genetic homogeneity of the Korean population, despite its diverse Y-chromosomal, mitochondrial, and HLA repertoires. This pattern reflects a long and continuous lineage history characterized by persistent internal admixture and genomic homogenization over thousands of years on the Korean peninsula. We also identified 16.8 million genomic variants that directly modify CG sites by creating or abolishing CG dinucleotides, providing the population-scale evidence of coordinated genomic-epigenomic regulatory mechanism in Koreans.

## Introduction

Population-scale whole-genome sequencing (WGS) has enabled high-resolution investigations into global genetic diversity, population structure, and disease susceptibility. Initiatives such as the 1,000 Genomes Project (1KGP) and the Genome Aggregation Database (gnomAD) (1) have collectively established a foundation for cataloging genetic variants across global populations. More recent large-scale biobank projects, including the UK Biobank (2) and FinnGen (3), have further integrated genomic data with extensive phenotypic records, offering powerful platforms for genotype–phenotype association and precision medicine. Across Asia, a growing number of national and regional genome programs have begun to capture both local and continental genetic diversity. Notable projects include the Taiwan Biobank (4), Biobank Japan (5), and Singapore’s PRECISE-SG100K project (6, 7), which collectively have advanced genomic medicine in East and Southeast Asia.

Within Korea, whole-genome sequencing projects have progressed, beginning with Korea1K (2020) with 1,094 whole genomes under the Korean Genome Project (KGP) (8) and Korean Variant Archive 2 (KOVA2; 2022) with 5,305 genomes and exomes which partly includes KGP data (9), followed by Korea4K (2024) with 4,157 high-depth whole genomes (10), revealing Korean-specific genetic variants and diverse genotype-phenotype associations and establishing critical population references for East Asian genomics.

Despite these advances, no single resource to date has conclusively captured the full genetic spectrum of Koreans. Prior studies were limited by cohort size, sequencing depth, or lack of integrated omic and clinical data. The absence of a unified, deeply sequenced, and phenotypically annotated Korean genomic reference has hindered both population genetics and translational research.

Here, we present Korea10K, the most comprehensive genomic and multiomic resource of the Korean population to date, encompassing 10,239 high-depth whole genomes coupled with partially matched transcriptomic, epigenomic, and phenotypic datasets. This resource marks a new phase of Korean population genomics.

## Results

### Variant Discovery and Population Reference Imputation Panel for Koreans

WGS of 10,239 Korean individuals yielded the most comprehensive catalog of genomic variation in this population to date. We identified 61,223,704 autosomal variants, comprising 57,184,188 single-nucleotide variants (SNVs) and 4,039,516 short insertions and deletions (indels) from 9,000 unrelated Korean samples. This reduction from 10,239 to 9,000 in sample number reflects stringent quality control, kinship filtering, and removal of ancestral outliers, to ensure unbiased variant discovery and accurate population-scale allele frequency estimation. Among the identified variants, 9,880,066 (16.1%) were novel and absent from dbSNP (v157), which already incorporated variants from our earlier Korea4K project (derived from 3,617 unrelated Korean genomes). Low-frequency variants dominated these previously unseen variations, with 33.6% classified as singletons (AC = 1), 14.9% as doubletons (AC = 2), and 3.4% as ultra-rare (AC > 2, AF ≤ 0.0005), revealing previously uncharacterized variation within the Korean population (Fig. 1A). Across all frequency categories, most of the novel variants were classified as SNVs, except for the very rare ones, where 34.9% were small indels (22.1% deletions and 12.8% insertions; Fig. S1).

**Figure 1.**
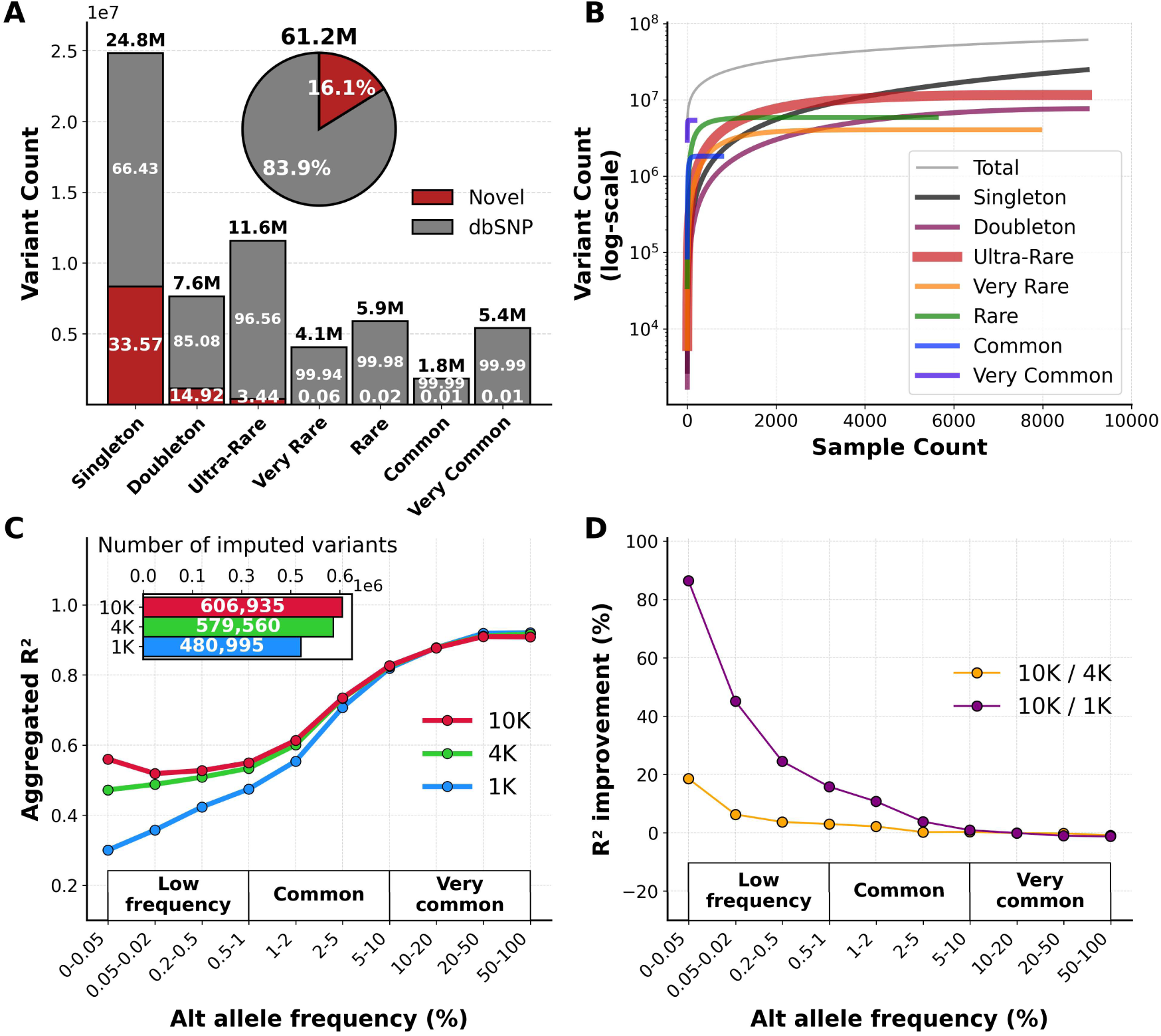
Variome landscape of the Korea10K whole genomes. **(A)** Stacked bar charts highlighting the distribution of discovered variant counts (y-axis) by allele frequency categories (x-axis), partitioned into novel (red) and dbSNP-reported (grey) variants. The pie chart summarizes overall variant statistics. **(B)** Cumulative variant discovery curve showing the number of variants identified with increasing sample size (y-axis, in log-scale) across allele frequency categories (x-axis). **(C)** Line plot depicting imputation accuracies (y-axis, as aggregated R^2^) across alternate allele frequency bins (x-axis) comparing Korea1K (blue), Korea4K (green), and Korea10K (red) population genome reference panels. The inset horizontal bar plot shows the total number of imputed variants (x-axis) per population genome reference panel (y-axis). **(D)** Line plot showing R^2^ improvement (y-axis, as percentage) comparing Korea10K to Korea1K (purple) and Korea10K to Korea4K (yellow) across allele frequency bins (x-axis). The allele frequency categories are defined as follows: Singleton (AC = 1), Doubleton (AC=2), Ultra-Rare (AC > 2, AF ≤ 0.0005), Very Rare (AF > 0.0005, AF ≤ 0.001), Rare (AF > 0.001, AF ≤ 0.01), Common (AF > 0.01, AF ≤ 0.05), and Very Common (AF > 0.05). Abbreviations: Alt allele = Alternative allele, AC = allele count, AF = allele frequency.

Cumulative variant discovery curves showed a continuous increase in novel variant counts with expanding sample size (Fig. 1B), including singletons and doubletons. Saturation analysis indicated that ultra-rare variants (AC > 2 and AF ≤ 0.0005) reached a mean discovery plateau at 8,955 samples (95% CI: 8,951-8,958), while very rare (AF > 0.0005 and AF ≤ 0.001) and rare variants (AF > 0.001 and ≤ 0.01) required 7,083 samples (95% CI: 7,026-7,141) and 4,692 samples (95% CI: 4,636-4,747), respectively. Common variants (AF > 0.01 and AF ≤ 0.05) saturated at mere 569 samples (95% CI: 557-581) while very common variants (AF > 0.05) saturated at 109 samples (95% CI: 106-111) (Fig. S2; Table S1). These results underscore the importance of large-scale sequencing to capture low-frequency Korean variation and justify the sample size of Korea10K for near-complete variant discovery for the traditional ethnic Korean population.

The Korea10K population reference panel demonstrated markedly improved imputation performance compared with earlier Korean panels (Korea1K and Korea4K), particularly for low-frequency variants (MAF < 1%; Fig. 1C). Relative to Korea1K, imputation accuracy increased by 15.8% to 86.4%, and by 3.0% to 18.6% compared with Korea4K in the descending order of allele frequency bins (table S2). The enhancement was most striking for ultra-rare variants, including singletons and doubletons (Fig. 1D). Furthermore, the Korea10K panel imputed 606,935 variants, representing a 26% increase over Korea1K (Fig. 1C, upper left), underscoring the value of progressive variant discovery for genotype imputation.

Korea10K provides a population reference of 61.2 million fully phased, functionally annotated autosomal variants, including 9.9 million novel alleles. Combined with a high-resolution Korean imputation panel, it enables accurate genotype imputation, rare-variant analyses, and population-specific functional interpretation.

### Population Structure and Genetic Homogeneity of Koreans

Principal component analysis (PCA) of autosomal genotypes positioned Koreans as a cluster within the East Asian genetic continuum, clearly separated from Han Chinese (CHB, CHS) and Japanese (JPT) populations from the 1KGP (Fig. 2A, B). Ethnically relevant reference genomes, KOREF1 (11) and its successor KOREF2, co-localized within the Korean cluster, validating them as well-suited genomic representatives. Admixture analysis revealed a single dominant ancestry component when modeling global autosomal variation with seven ancestral populations (K = 7), which best explained global autosomal genetic structure (Fig. 2C, top panel; Fig. S3). We additionally examined K = 9, which showed the second-lowest cross-validation error, to explore finer-scale population structure within East Asia. At higher cluster numbers, Koreans exhibited a dual-ancestry pattern that was also observed in northern Han Chinese and Japanese populations. Notably, this pattern persisted even among 485 individuals selected based on genetic distance to maximize within-population diversity (Fig. 2C, bottom panel), reflecting long-standing demographic continuity across the Korean Peninsula.

**Figure 2.**
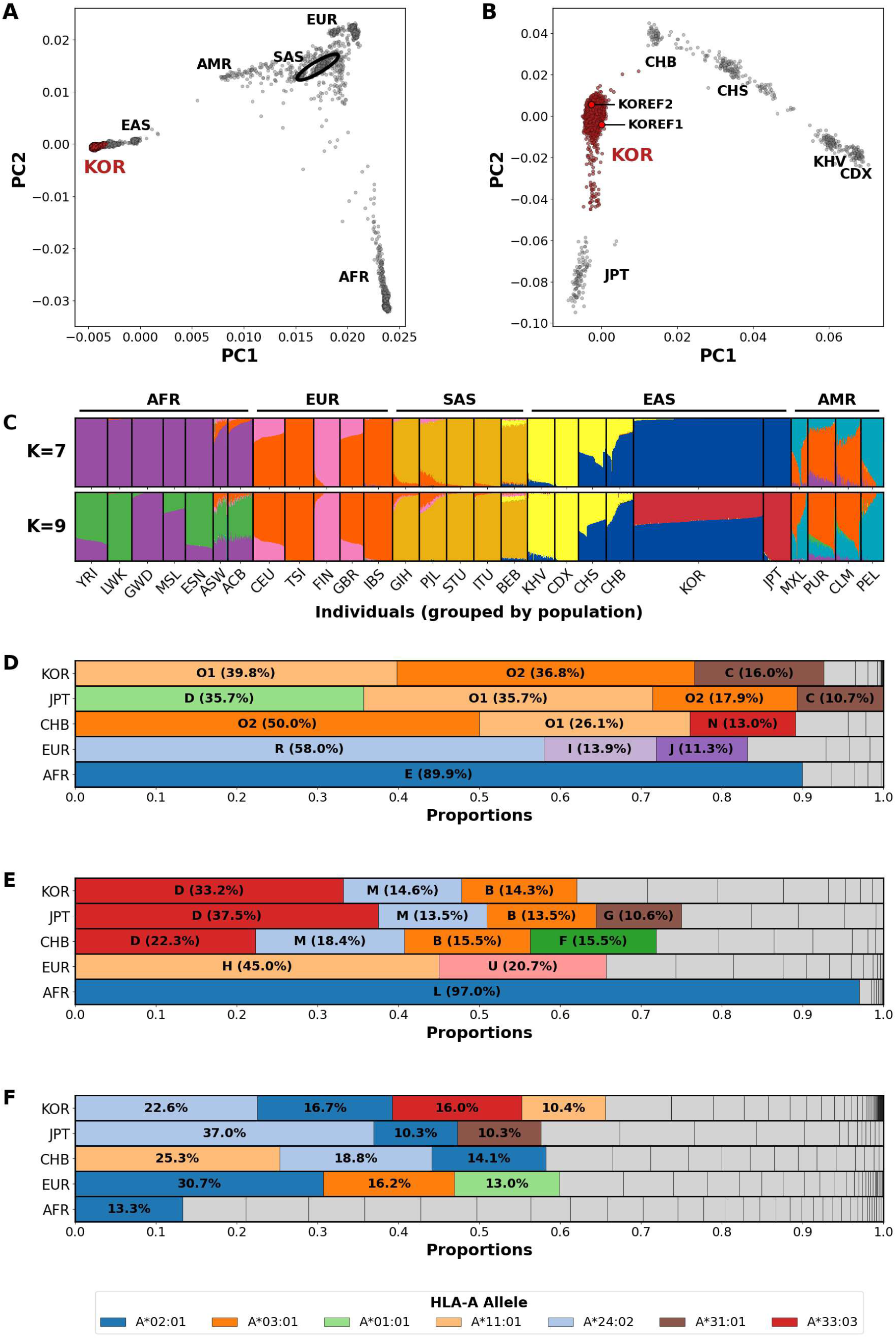
Population structure and uniparental lineage composition of the Korean population. **(A)** Global principal component (PC) bi-plot of PC1 (x-axis) and PC2 (y-axis) based on autosomal variants from 9,000 unrelated Koreans (KOR) and 2,541 individuals from the 1000 Genomes Project (1KGP). Koreans are displayed as closed red dots. SAS placed within AMR are open-circled with a black ellipse for distinction. **(B)** East Asian PC bi-plot of PC1 (x-axis) and PC2 (y-axis) showing 9,000 unrelated Koreans (KOR) and 501 samples in other East Asian populations (CHB, CHS, CDX, JPT, and KHV) of 1KGP. Koreans are displayed as closed dim red dots. The Korean References (KOREF1 and KOREF2) are denoted as bright red dots. **(C)** Admixture plots at K=7 (top panel) and K=9 (bottom panel). The x-axis displays individuals in analysis that have been grouped by population. The y-axis denotes number of admixture clusters in analysis (i.e., K). **(D)** Stacked bar chart showing proportions (x-axis) of major Y-chromosomal (except O1 and O2 for scrutiny) haplogroups of 5,290 unrelated Korean males compared to other populations (y-axis). **(E)** Stacked bar chart showing proportions (x-axis) of major Mitochondrial (MT or mt) haplogroups of 9,000 unrelated Koreans and other populations (y-axis). **(F)** Stacked bar chart showing proportions (x-axis) of 18,000 HLA-A alleles of 9,000 unrelated Koreans compared to other populations (y-axis). Abbreviations: AFR = African, EUR = European, SAS = South Asian, EAS = East Asian, AMR = Ad Mixed American, KHV = Vietnamese, CDX = Chinese Dai, CHS = Southern Han Chinese, CHB = Northern Han Chinese, KOR = Korean in Korea10K, JPT = Japanese in Tokyo, KOREF = Korean Reference, 1KGP = the 1000 genome project.

Analysis of 5,290 unrelated Korean Y-chromosomal genomes revealed a strong dominance of haplogroup O (N = 4,053, 75.7%), comprising both O1 (N = 2,105, 39.8%) and O2 (N = 1,948, 36.8%), followed by C (N = 847, 16.0%). This distribution is broadly consistent with patterns observed across northern East Asia (Fig. 2D). In contrast, population-specific signatures include markedly higher frequencies of haplogroup D in the Japanese (35.7%) and a stronger representation of O2 in northern Han Chinese (50.1%). Outside East Asia, Europeans are dominated by haplogroup R (58%), whereas Africans show overwhelming prevalence of haplogroup E (89.9%), illustrating broad continental divergence in male ancestries. Phylogenetic reconstruction based on pairwise Y-chromosomal genetic distances from 5,871 haplogroup-classified males showed limited resolution of major haplogroups and their subclades. The limited phylogenetic resolution reflects shallow sequence divergence among lineages due to recent demographic expansions, probably since bronze age population explosion (12), into the Korean peninsula and the persistence of shared ancestral polymorphisms (Fig. S4A).

Mitochondrial genomes in Koreans exhibited markedly greater diversity compared to the Y chromosome. A total of 10,239 mitochondrial assemblies and alignments were generated, representing one of the largest population-level mitogenomic resources to date. Phylogenetic reconstruction with the complete circular assemblies revealed a deeply branching topology, indicating highly diverse maternal lineages in Korea (Fig. S4B). Despite extensive overall diversity, mitochondrial lineages in Koreans were primarily partitioned into two major macrohaplogroups, M and N, encompassing nearly all dominant subclades, namely D (N = 2,988, 33.2%), M (N = 1,315, 14.6%), and B (N = 1,285, 14.3%) in 9,000 unrelated Koreans (Fig. 2E). The absence of Western Eurasian (H and X) and African haplogroups (L) supports prolonged maternal lineage isolation due to Korea’s geographic remoteness (Fig. S4B). PCA of 1,247 mitochondrial variants further captured fine-scale maternal sub-structure concordant with haplogroup-defined clusters (Fig. S5). These findings support multiple migration and admixture events involving females from neighboring East Asian populations into the Korean peninsula (13).

At the immunogenetic level, HLA class I (HLA-A) allele frequencies illustrate characteristic features of the Korean population. Among 9,000 unrelated Koreans (18,000 haplotypes), four alleles, including A*24:02 (N = 4,059, 22.6%), A*02:01 (N = 3,003, 16.7%), A*33:03 (N = 2,879, 16.0%), and A*11:01 (N = 1,871, 10.4%), collectively accounted for nearly two-thirds of all haplotypes (Fig. 2F), indicating a narrow and coherent allelic spectrum. In comparison, Japanese individuals show a stronger enrichment of A*24:02 (37.0%), whereas Han Chinese exhibit a predominance of A*11:01 (25.3%); both populations display reduced representation of A*33:03. By contrast, Europeans and Africans exhibit markedly broader HLA polymorphism, with A*02:01 comprising 30.7% of European haplotypes but accompanied by numerous low-frequency variants, and no single allele exceeding 13% in African populations.

Collectively, Koreans represent a genetically homogeneous population shaped by long-term geographic isolation and demographic continuity, with additional genetic contributions from multiple prehistoric admixture events involving both male and female migrants from neighboring East Asian populations. The ancestral and immunogenetic resources generated by Korea10K offer a valuable foundation for population genetics, forensic analyses, and immunological research.

### Population-Specific CG Context-associated Variants (CGV) in Koreans

DNA mutations can directly alter CG dinucleotide contexts, which may in turn modulate DNA methylation potential and downstream gene regulation. Although direct methylation data were not analyzed here, we profiled sequence-driven changes in CG context-associated variants (known as CGVs, CG-SNPs, or meSNPs), a form of geno-epigenetic polymorphism. These CGVs suggest population-level differences in epigenetic regulation and gene expression, particularly in pathways related to psychiatric and metabolic disease risk in the Korean population.

We performed a whole-genome analysis of 9,000 unrelated Korean individuals and identified extensive variation affecting the CG dinucleotide context. This provides a population-scale view of CG sequence diversity. Among 61,223,704 high-quality biallelic variants, 16,388,405 (26.8%) directly altered CG sites, comprising 7,154,078 (11.7%) CG-creating and 9,364,566 (15.3%) CG-eliminating variants (Fig. 3A, B). The proportion of CG-altering variants was consistent across allele frequency categories, indicating that CG context changes occur broadly across the frequency spectrum (Fig. S6).

**Figure 3.**
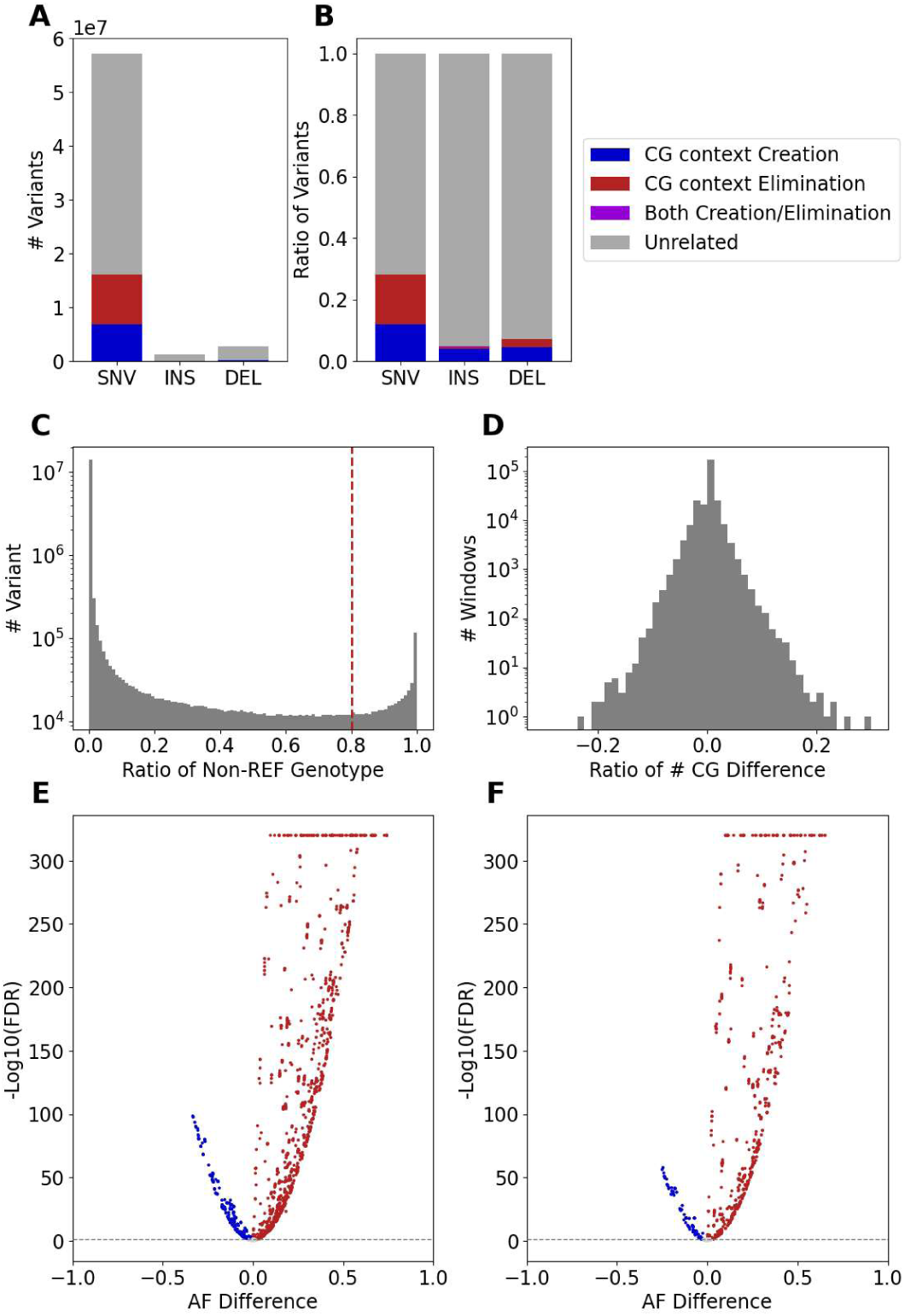
Population-specific CG context-associated variants (CGVs) in Koreans. **(A)** Total number (y-axis) of single nucleotide variants (SNVs), insertions (INS), and deletions (DEL) (x-axis) affecting CG dinucleotide context. **(B)** Proportion (y-axis) of different variant types (x-axis) categorized by CG context impact: creation (blue), elimination (red), both creation/elimination (purple), and unrelated (gray). **(C)** Distribution of variant counts (y-axis) over the ratio of non-reference genotypes (x-axis) across all individuals (dashed line = Korean-popular threshold, 0.8). **(D)** Relationship between the number (y-axis) and ratio (x-axis) of CG site differences per genomic window. **(E-F)** Volcano plots showing allele frequency (AF) differences between Koreans and Europeans (x-axis) for CG-creating and CG-eliminating variants, respectively, highlighting population-specific enrichment (y-axis, as negative of log_10_FDR).

At the population level, 416,796 variants were both CG context-related and common in Koreans (allele frequency ≥0.8; Fig. 3C), including 212,621 CG-creating and 206,845 CG-eliminating variants. A smaller subset (108,824 variants; 0.17%) simultaneously created and eliminated CG motifs through adjacent substitutions or small indels, leading to positional motif shifts without altering total CG abundance, an underappreciated source of bias in methylation-based analyses.

Regional aggregation of CGVs in 10-kb windows revealed that 97.6% of the genome differed by <5% in CG density relative to GRCh38 (Fig. S7), whereas 408 regions (0.15%) exhibited >10% deviation (266 increases, 142 decreases; Fig. 3D). The top five CG-altering “hotspots” overlapped functionally important loci, including major histocompatibility complex (MHC) and leptin receptor (LEPR) regions, both key regulators of immune and metabolic homeostasis (14). Notably, LEPR exhibited a 23.1% relative CG gain, ranking among the top loci (Table S3), suggesting potential methylation-mediated modulation of expression.

Comparison with high-depth 1KGP European genomes (N = 522) showed that 72.8% of the CGVs in the hotspot regions (with >10% deviation against GRCh38) were significantly more frequent in Koreans (2,078/2,853 variants; CG-creating: 1,320/1,803; CG-eliminating: 758/1,050; Fig. 3E, F). Genes within CG-gain regions were enriched for functions related to neuropsychiatric disorders, synaptic signaling, and metabolic regulation (Fig. S8, 9).

Together, these findings establish a catalog of Korean-specific CG context-associated variants (CGVs) that reshape the CG landscape at the population level. This resource supports variant-aware epigenetic modeling, enhances population-specific methylation probe design, and provides a foundation for interpreting genetic-epigenetic interactions underlying disease risk and regulatory evolution in East Asians.

### Korea10K as a Genomic, Multiomic, and Phenomic Data Resource

The Korea10K project expands upon earlier phases including KPGP (N = 118), Korea1K (N = 1,094), and Korea4K (N = 4,157). Korea10K established a nationwide, population-scale genomic and multiomic resource comprising 10,239 high-depth WGS from Korean individuals (Fig. 4A). Participants were recruited through multiple regional centers and institutional partners across sequential collection phases, achieving broad demographic representation and phenotypic diversity.

**Figure 4.**
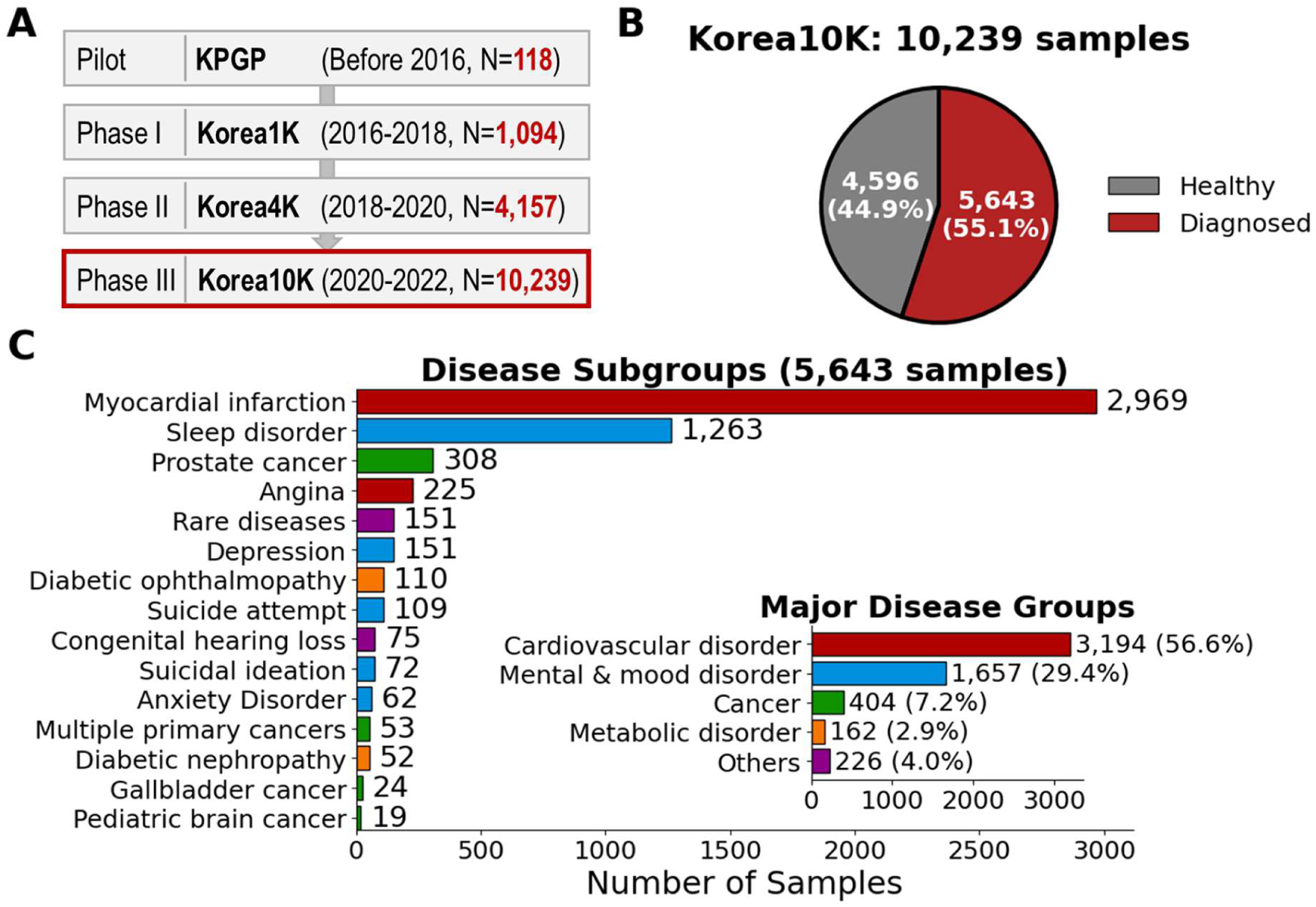
Overview of the Korea10K cohort. **(A)** Sequential recruitment phases of the Korea10K project, expanding from a pilot study (KPGP) to large-scale national inclusion through Korea1K, Korea4K, and Korea10K, comprising 10,239 high-depth whole genomes collected in total. **(B)** Distribution of healthy (N = 4,596) and clinically diagnosed (N = 5,643) participants. **(C)** Composition of disease subgroups and major disease categories (y-axis) among diagnosed participants (x-axis). Cardiovascular disorders represented the largest group (N = 3,194), followed by mental and mood disorders (N = 1,657), cancers (N = 351), metabolic disorders (N = 162), others (N = 226), and. Abbreviation: KPGP = Korean Personal Genome Project.

Among all participants, 44.9% (N = 4,596) were healthy volunteers without apparent disease onset at enrollment, whereas 55.1% (N = 5,643) had clinically verified diagnoses (Fig. 4B). Disease sub-cohorts encompassed cardiovascular, neuropsychiatric, metabolic, oncologic, and congenital conditions. Cardiovascular diseases represented the largest group (56.6%), including acute myocardial infarction (AMI) and angina, followed by mental and mood disorders such as depression and sleep disorders (29.4%), with additional subgroups to support disease-specific analyses (Fig. 4C).

Korea10K marks a major expansion of the KGP into the multiomic era. Population-scale molecular profiles from blood include 868 transcriptomes, 1,211 DNA methylomes, and 102 chromatin accessibility datasets, and six proteomes, providing a systems-level view of regulatory effects of genetic variation and mechanistic insights into phenotypes. Complementary phenotypic resources include the Health Check-up (HC) dataset (6,583 participants, 336 phenotypes; table S4) and the Lifestyle Questionnaire (LQ) dataset (4,923 participants, 370 survey measures; table S5), encompassing diet, physical activity, smoking, alcohol use, sleep, and psychosocial factors (Table 1).

**Table 1.**
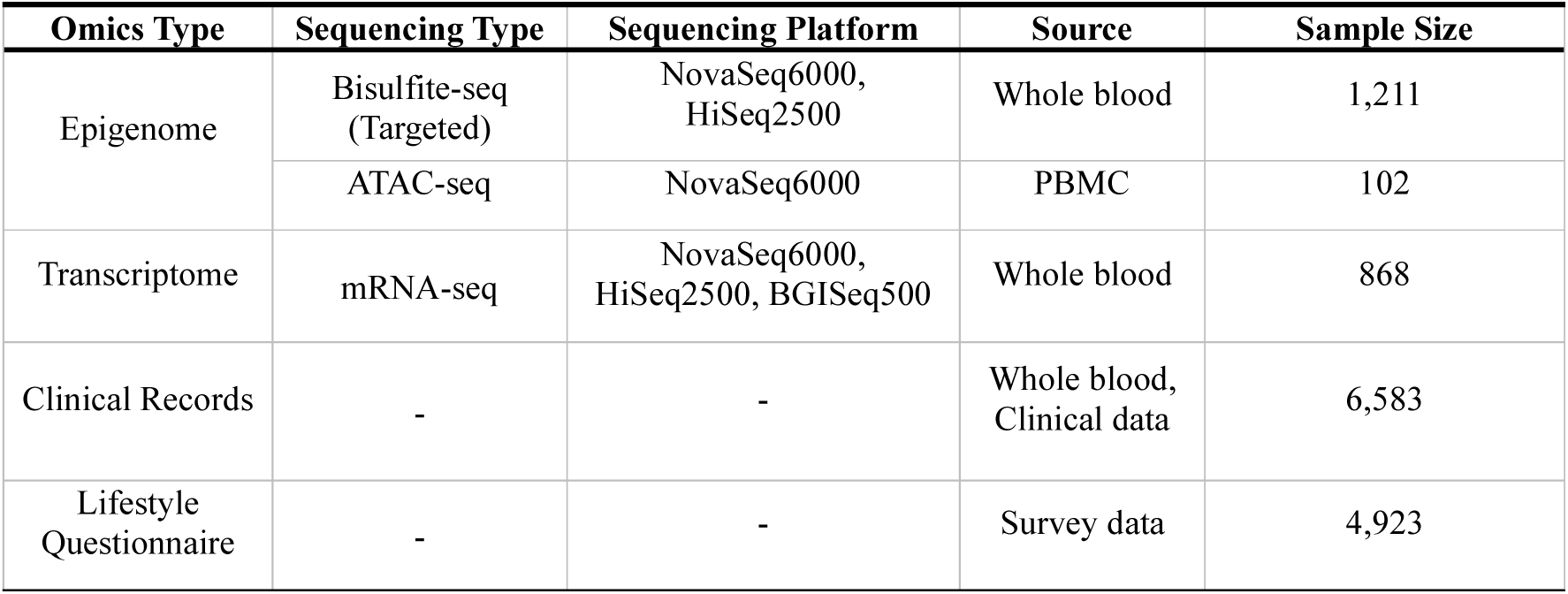
Overview of Multiomic and Phenomic Data Resource in Korea10K. PBMC = Peripheral Blood Mononuclear Cell.

KOREF (KOrean REFerence) represents one of the most precise longitudinal omics profiling resources available in KGP. KOREF1 (middle-aged donor, PGP ID of hu3D760A) and KOREF2 (younger donor) provide high-precision datasets, including 39 WGS, 36 epigenomes (6 Hi-C, 19 DNA methylomes, 11 ATAC-seq), 73 transcriptomes, 6 proteomes (SomaScan assays), six microbiomes (four 16S rRNA-seq, two metagenomic WGS), and 12 phenomes (ten clinical records and two lifestyle information datasets), collected over a decade and a half-decade, respectively. Also, KOREF1 is the first Korean multiomic reference accompanied by the first fully phased diploid Korean genome, achieving high-accuracy and gapless telomere-to-telomere completeness (15). KOREF2 constitutes the next-generation reference with a rich set of third-generation sequencing data (PacBio HiFi and ultra-long Oxford Nanopore data) toward near-complete assembly, improving the mappability of the multiomic data for highly accurate reference-based analyses.

Collectively, Korea10K provides a resource capturing population-scale genomic and multiomic variation representative of the Korean population including reference-quality longitudinal omics datasets.

### Genetic Associates of Clinically-Inferred Biological Aging in the Korea10K Cohort

To demonstrate the utility of the Korea10K data resource, we performed a genome-wide association analysis of biological aging, quantified as Phenotypic Age Advance (PAA) (16), among 4,498 unrelated Koreans in Korea10K with HC data available. PAA was defined as the residual of phenotypically measured biological age (PhenoAge) regressed on chronological age.

PhenoAge was calculated from clinical biomarkers associated with 10-year cumulative all-cause mortality risk. Among 12 candidate biomarkers available in both Korean and U.S. reference cohorts, a mortality model trained in the Third National Health and Nutrition Examination Survey dataset (NHANES III; N = 11,562) identified four biomarkers with biologically meaningful effect sizes (HR > 1.000 at 4 s.f.): serum creatinine, chronological age, systolic blood pressure, and blood glucose (Fig. 5A). Their hazard ratios were 1.338 (95% CI: 1.154-1.552), 1.051 (95% CI: 1.049-1.053), 1.009 (95% CI: 1.007-1.011), and 1.001 (95% CI: 0.999-1.002), respectively, yielding a concordance index of 0.885. The composite partial hazard score constructed from these variables significantly discerned deceased from surviving individuals in both the discovery (NHANES III) and validation (NHANES IV) cohorts (P < 10-5; Fig. 5B, C). A multivariate Gompertz model parameterized with these biomarkers was subsequently calibrated for Korean clinical profiles to compute PhenoAge and derive PAA (Fig. S10).

**Figure 5.**
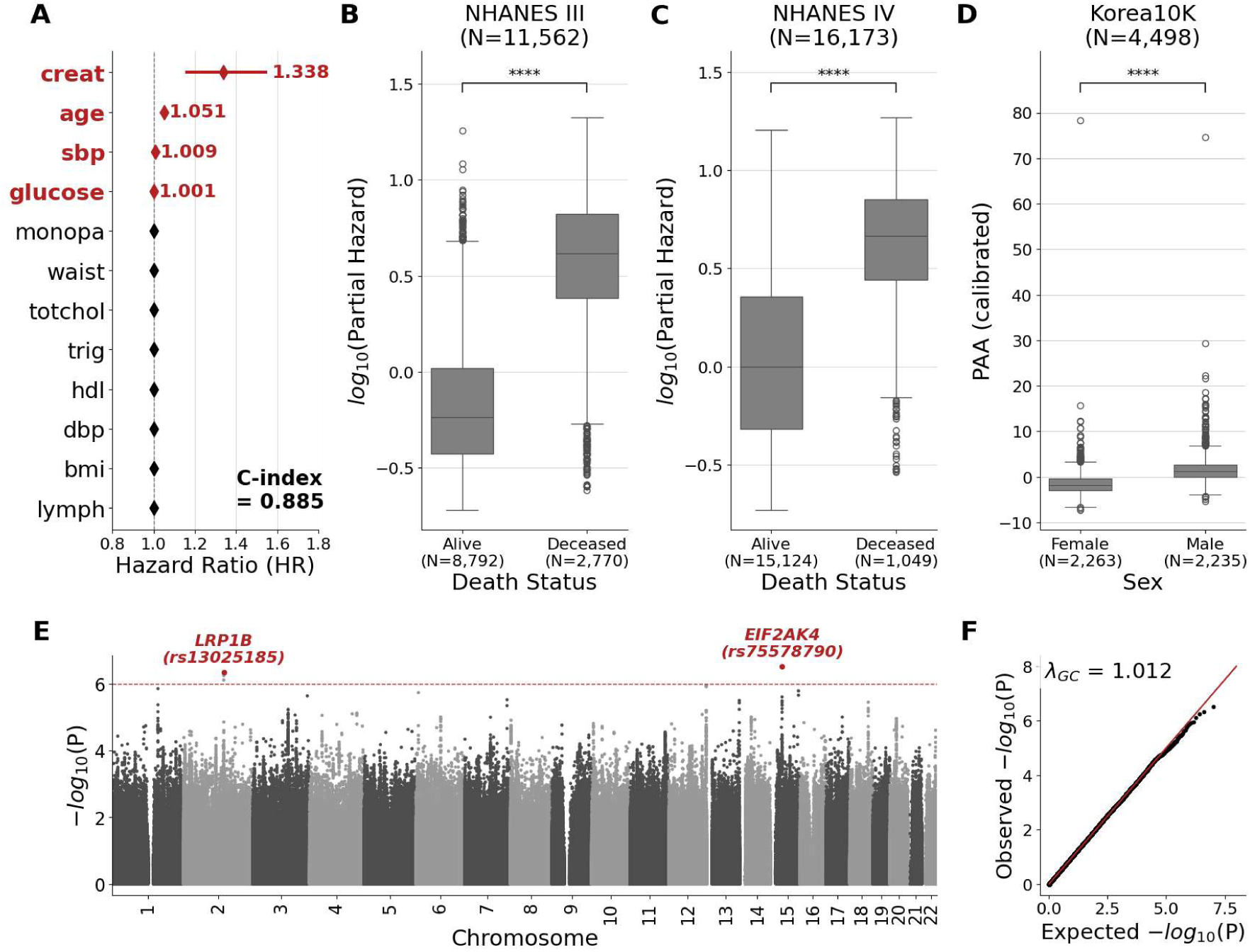
Clinical and genetic correlates of biological aging. **(A)** Forest plot of a multivariable Cox proportional hazards model for 10-year all-cause mortality in NHANES III, displaying hazard ratios as closed diamonds and 95% confidence intervals as error bars (x-axis) for clinical covariates (y-axis); red markers indicate variables with biologically meaningful effect sizes, and the concordance index (C-index) of the model is shown at the lower right. **(B, C)** Box plots of the composite mortality risk score (log_10_ Partial Hazard; y-axis), stratified by death status (x-axis) in the training NHANES III cohort (B) and an independent test NHANES IV cohort (C). Boxes denote the interquartile range (IQR), center lines the median, whiskers 1.5× the IQR, and point outliers (open circles), with group differences assessed using two-sided Mann-Whitney U tests. **(D)** Box plots of calibrated phenotypic age acceleration (PAA; y-axis) in the Korea10K cohort, stratified by sex (x-axis), with group comparison evaluated by a two-sided Mann-Whitney U test. **(E)** Manhattan plot of genome-wide association analysis of PAA-INT, displaying significance of each variant (y-axis) across genomic position (x-axis). The red dashed line marks the suggestive significance threshold (*P* < 10^-6^); loci and lead variants exceeding this threshold are highlighted in red. **(F)** Quantile–quantile (QQ) plot of observed (y-axis) versus expected (x-axis) −log_10_(*P*) values for the GWAS in panel (E), with the genomic inflation factor (λ_GC_) shown at upper left. Abbreviations: age = chronological age, bmi = body mass index, creat = serum creatinine, dbp = diastolic blood pressure, GC = genomic control, glucose = blood glucose, GWAS = Genome-wide association study, hdl = high-denstiy lipoprotein, lymph = lymphocyte percentage, monopa = monocyte percentage, NHANES = U.S. National Health and Nutrition Examination Survey, sbp = systolic blood pressure, totchol = total cholesterol, trig = triglyceride, waist = waist circumference.

Among 4,498 unrelated Koreans, PAA exhibited a right-skewed distribution and differed significantly by sex (P < 10-5; Fig. 5D); sex was excluded as a covariate to avoid adjusting away biologically relevant variation. To mitigate the influence of skewness and outliers and ensure robust association testing, PAA values were inverse-normal transformed based on their ranks (i.e., PAA-INT).

We tested 6,996,863 autosomal variants for association with PAA-INT, identifying four SNVs at two unique loci (*LRP1B* and *EIF2AK4*) that surpassed the suggestive significance at P < 10-6 (Fig. 5E). Three SNVs (rs4954893, rs13016322, and rs13025185) were positioned on *LRP1B* locus, while one (rs75578790) mapped to *EIF2AK4* (Fig. 5D). LRP1B is implicated in cognitive performance for successful aging (17). EIF2AK4 (or GCN2) is a central regulator of the integrated stress response (18). Although the GWAS exhibited mild deflation, genomic inflation was minimal (λ_GC_ = 1.012; Fig. 5F), indicating effective control of confounding despite the modest sample size.

Together, these results highlight the capacity of Korea10K to uncover biologically plausible genetic contributors to aging and demonstrate its value as a foundational population-scale resource for dissecting the molecular mechanisms of health and diseases in East Asian populations.

## Discussion

There are several limitations to this study. From a design perspective, Korea10K is currently a cross-sectional resource except two KOREF samples; the absence of longitudinal outcome data precludes tracking of biological aging, disease progression, and causal inference over time. Although we classified participants in Korea10K as either healthy or clinically diagnosed, the putative healthy group cannot be assumed to be physiologically normal or free of undetected conditions, and diagnostic misclassification remains possible within the clinically diagnosed cohort despite clinician verification. More precise, data-driven stratification of participants into biologically informed subgroups will be critical for improving the resolution and interpretability of downstream association analyses (19). Clinical and lifestyle questionnaire data were collected only for a subset of individuals directly enrolled through participating institutions. Samples acquired via Material Transfer Agreements (MTAs) from external Korean biobanks lacked detailed phenotypic information. This incomplete metadata reduced the effective sample size for genome-wide association studies (GWAS), limiting power for large-scale genotype-phenotype mapping beyond the Korea4K dataset. Budgetary constraints restricted the generation of additional multiomic datasets, limiting systems-level interpretation of genetic effects. For instance, population-matched whole blood RNA-seq would enable transcriptome-wide association analyses (TWAS) and expression QTL (eQTL) mapping tailored to the Korean population, improving fine-mapping and gene prioritization (20). DNA methylation profiling and ATAC-seq would allow investigation of methylation QTLs (meQTLs) and chromatin-accessibility QTLs (caQTLs) facilitating genome-epigenome integration to reveal the core regulatory mechanisms (21, 22).

Technical limitations also remain. Reliance on short-read sequencing and an incomplete linear reference genome (GRCh38) left substantial genomic regions, including centromeres, sub-telomeres, segmental duplications, and the Y chromosome, poorly resolved. Consequently, structural variants, complex rearrangements, and tandem-repeat expansions were incompletely captured. Reference-based epigenomic analyses similarly remain constrained by reference bias. CGVs delineate regions where the Korean genomic CG landscape diverges from the GRCh38 human reference genome, limiting accurate characterization of population-specific methylation states and chromatin architecture. The prevalence of CGVs underscores the need for sequence-context aware modelling. Adoption of long-read sequencing (23), ethnically-specific (or even personal) assemblies such as KOREF1 (11), and graph-based pangenomes (24) will also be essential for achieving comprehensive variant representation and their accurate functional annotation.

Future expansions incorporating longitudinal phenotyping, long-read sequencing, and multiomic profiling will enable more precise capture of structural variation, population-specific epigenetic landscapes, and elucidation of molecular mechanisms underlying aging and disease. We expect that cross-platform multiomics signatures will elucidate the gene regulatory mechanisms underlying Korean-specific disease biomarkers through integrated pathway analysis.

Despite limitations, Korea10K substantially extends both the scale and depth of Korean population genomics. Compared with Korea4K with 4,157 individuals (3,617 unrelated, 45 million variants), the cohort expanded to 10,239 genomes (9,000 unrelated) identified 61 million variants, including nearly 10 million previously unseen alleles, approaching the saturation of ultra-rare alleles. This near-complete discovery of low-frequency and private variants improved imputation performance and fine-mapping resolution, thereby enhancing the statistical power of future GWAS and rare-variant association studies in Koreans.

Population structure analyses confirmed again that Koreans form a distinct cluster within East Asia. Autosomal PCA positions them along the East Asian continuum, adjacent to Japanese and northern Han Chinese populations, reflecting shared ancestry while preserving a unique genetic profile. Y-chromosomal haplogroups point to recent male-mediated expansions consistent with Neolithic agricultural dispersals (12). Big mitochondrial diversity preserves signatures of ancient female-mediated admixture and Bronze Age demographic expansions (13). Immunogenetic profiles also reveal a broadly shared dominance of HLA-A alleles across northern East Asia, with a selective enrichment of A*33:03 in Koreans. Population-specific alleles suggest local selection pressure from endemic pathogens, with implications for vaccine response (25), autoimmune disease susceptibility (26), and drug contraindications (27). The exceptional homogeneity of Koreans minimizes population stratification bias and enhances the power, precision, and reproducibility of genomic and multiomic association studies (28).

Korea10K has already facilitated the discovery of genomic and multiomic biomarkers associated with diverse phenotypes, including cardiovascular disorders (acute myocardial infarction) (29–32), neuropsychiatric traits (depression, anxiety, and suicidal behavior) (33, 34), infectious diseases (COVID-19) (35), and biological aging (36), providing mechanistic insights into disease etiology and progression.

By leveraging genetic homogeneity, high-resolution multiomic data, and deep phenotyping, Korea10K provides a robust platform for biomarker discovery, disease mechanism elucidation, and the development of precision medicine strategies in Korea.

## Materials and Methods

### Data Availability

Genetic variations data from Korea10K that pose no privacy concerns are available in VCF format through the Korea BioData Station (K-BDS, https://kbds.re.kr/), provided by the Korea Bioinformation Center (KOBIC). Access to raw sequencing data can be granted upon request and is subject to data-use agreements. Access to individual-level datasets additionally requires approval from the relevant Institutional Review Board (IRB) and Data Access Committee, in accordance with international guidelines for human genomic data sharing. The 1KGP Phase 3 high-depth (30×) dataset used in this paper was retrieved from the Amazon Web Services (AWS) server (https://registry.opendata.aws/1000-genomes). The 1KGP Phase 1 legacy data can be found on their FTP server (https://ftp.1000genomes.ebi.ac.uk/vol1/ftp/release/20130502/).

### Ethics, Consent, and Permissions

Sample collection and sequencing were approved by the Institutional Review Board (IRB) of the Ulsan National Institute of Science and Technology (UNISTIRB-15-19-A and UNISTIRB-16-13-C). The data employed in our study originates from voluntary blood or saliva donations, and we have diligently secured explicit, comprehensive consent forms from the participants prior to sample collection. These consent forms explicitly outline the intended use of their data for research purposes and underscore the voluntary nature of their participation. Our study adheres to the ethical guidelines and regulations stipulated by the IRB. As a result, we can make the data available under controlled access while respecting the privacy and consent of the participants. Any data set we provide can be subject to deletion in the future according to policy guidelines given by national and institutional authorities and regulations.

### Sample Collection

Peripheral blood or saliva samples were collected using standardized procedures. Genomic DNA was extracted using the DNeasy Blood & Tissue Kit (Qiagen) for blood and the GeneAll Exgene Clinic SV Mini Kit for saliva, following the manufacturers’ protocols. DNA quality and quantity were assessed using Qubit 3.0 Fluorometer (Thermo Fisher Scientific) and Bioanalyzer 2100 (Agilent) for library preparation.

### Whole-Genome Sequencing

Whole-genome sequencing (WGS) was performed at a mean depth of 30× using MGI DNBSEQ-T7 (N = 6,311; 61.6%), Illumina NovaSeq 6000 (N = 2,832; 27.7%), and HiSeq X Ten (N = 1,096; 10.7%) (Fig. S11; Table S6). Early-phase sequencing employed Illumina platforms, whereas large-scale production was performed on the MGI T7 platform for high-throughput efficiency. High-molecular-weight DNA was enzymatically fragmented, and libraries prepared using platform-specific kits: TruSeq Nano or TruSeq PCR-Free (Illumina) and MGIEasy FS DNA Library Prep (BGI), followed by circularization and rolling circle amplification for DNBSEQ-T7 libraries. Libraries were sequenced as 2 × 150bp (or 151bp for Illumina) paired-end reads.

### Read Processing and Alignment

Raw sequencing reads were trimmed with fastp v0.23.2 (35) to remove platform-specific adapters, polyG tails, and reads with low-quality bases (mean base quality below Q20). Trimmed reads were aligned to the GRCh38 reference genome using BWA-MEM v0.7.17 (36) with read group information. Alignments were coordinate-sorted and duplicates marked with Picard v2.27.5 (37). Base quality score recalibration was performed using GATK BaseRecalibrator and ApplyBQSR v4.6.1.0 (38).

### Variant Calling and Joint Genotyping

Per-sample variant calling was performed using GATK HaplotypeCaller v4.6.1.0 in GVCF mode, adjusting ploidy for sex chromosomes. GVCFs from 10,239 samples were aggregated using GenomicsDBImport and jointly genotyped with GATK v4.6.1.0. Post-calling, variant quality score recalibration (VQSR) was applied separately to SNPs and indels using dbSNP v156, Mills & 1000G Gold Standard, HapMap, and Omni arrays. Final VCFs were compressed, indexed, and standardized according to VCF v4.2 specifications.

### Sample and Variant Filtering

Joint-called variant call format (VCF) files from 10,239 Korean individuals were subjected to multi-step quality control (QC) to obtain a high-quality, unrelated cohort of 9,000 individuals for downstream analyses. Autosomal and X-chromosomal (chrX) VCFs were first filtered to retain biallelic variants that passed variant quality score recalibration (VQSR). For Y-chromosomal (chrY) VCFs, the VQSR was not applied. All VCFs were converted to PLINK2 format for downstream analyses.

At the sample level, individuals with genotype missingness exceeding 10% or heterozygosity beyond ±3 standard deviations (SD) from the mean were excluded (205 samples). Heterozygous allele balance (ABHet) was examined to identify samples with excessive allelic imbalance; individuals with mean ABHet outside ±2 SD were removed (221 samples). Relatedness was assessed using KING from PLINK v2.0.0-a.6.13LM (39), and one individual from each pair with kinship coefficients > 0.125 (third-degree relatives or closer) was removed (800 samples). Population analysis using principal component analysis of 9,013 “supposed” Koreans with 1000 Genomes Project (1KGP) variome detected non-Korean outliers (13 samples; Methods: Population Structure and Admixture Analyses with EBI’s 1KGP Reference).

Variant-level QC was applied to remove variants with: (i) ABHet outside 0.4–0.6, (ii) allele balance in homozygous genotypes >10%, (iii) >10% of genotypes with zero allele depth, (iv) ExcessHet > 60, (v) call rate < 99%, or (vi) Hardy-Weinberg equilibrium (HWE) p-value < 1×10⁻⁶. After applying all filtering steps, 61,223,704 high-confidence biallelic variants were retained across 9,000 independent Korean genomes for downstream analyses. The variants were annotated using Variants Effect Predictor (VEP) v115 (40).

### Variant Discovery Saturation Analysis

To assess the saturation of variant discovery, all 9,000 unrelated individuals were included in each analysis, and the cumulative number of unique variants was recorded as samples were sequentially added. Because cohort heterogeneity, including population substructure and disease-specific variant profiles, can bias saturation estimates depending on the order in which samples are included, we performed 100 independent permutations in which the sample order was randomly shuffled without replacement. For each permutation, the cumulative variant discovery curve was generated, and saturation was defined as the median point at which all unique variants had been observed across permutations. This permutation-based approach ensures that the saturation estimate is robust to cohort composition and sample ordering, rather than reflecting artifacts introduced by the inclusion of individuals with atypically high private variant loads or shared genetic backgrounds.

### Population Reference Panel Construction and Benchmarking Imputation Performance

A Korean-specific population reference panel was constructed from 9,013 unrelated genomes. Autosomal variants were phased using SHAPEIT5 v5.1.1 (commit = 990ed0d) (41), and imputation was performed with Minimac3 v2.0.1 (42). Accuracy was evaluated as the squared Pearson correlation (R²) between imputed dosages and observed genotypes, stratified by allele frequency using the “GetCorrelation” from the TOPMed Imputation evaluation pipeline (https://github.com/sgagliano/UKB_WES_vs_TOPMed_IMP) (43). For variants exhibiting a negative correlation, the negative sign was retained in the squared correlation value to preserve directional information. Korea10K (this study), Korea4K, and Korea1K reference panels were tested their imputation performance against same dataset as in the previous study (7).

### Population Structure and Admixture Analyses with EBI’s 1KGP Reference

A total of 3,202 high-coverage (30×) whole-genome datasets from the 1000 Genomes Project (1KGP) were retrieved (44). After applying the same kinship-filtering criteria used for Korea10K (Methods: Sample and Variant Filtering), 2,541 unrelated individuals were retained. These samples were jointly merged with 9,013 putative Korean genomes. Variant harmonization was performed using reciprocal intersection of SNPs present in both datasets PLINK2 (--extract-intersect), ensuring positional and allelic concordance prior to merging. The two VCFs were then merged using PLINK v1.9.0-b.7.11 (--bmerge) (45), producing a unified dataset containing 15,501,578 unfiltered autosomal variants.

Quality control was applied uniformly across all samples. Variants were excluded if they met any of the following thresholds: (i) minor allele frequency (MAF) < 1%, (ii) deviation from Hardy-Weinberg equilibrium (HWE) with P < 10^-6^, and (iii) missing genotype rate > 1%. After filtering, 4,315,937 variants were retained. To reduce redundancy due to linkage disequilibrium (LD), variants were LD-pruned using a 200 kb sliding window with the parameters --indep-pairwise 200kb 1 0.5, resulting in 850,281 variants for population structure analysis.

Merging the 9,013 Korean genomes with all 2,541 1KGP individuals yielded 850,871 LD-pruned variants shared across all populations. When restricting the merge to the 501 East Asian (EAS) individuals in 1KGP, 876,547 QC-passed and LD-pruned variants were obtained and used for principal component (PC) computation.

For model-based ancestry estimation, ADMIXTURE v1.3.0 (46) was run for from K = 1 to K = 17. To reduce computational burden and account for long-range LD, variants were further pruned using --indep-pairwise 500kb 1 0.2. Identity-by-state clustering was performed using PLINK1 (--cluster --K 5), producing 500 genetic clusters. Clusters containing ≤ 2 samples were removed, leaving 485 clusters. For each cluster, PCA was performed on the first five components (PC1–PC5), and the sample closest to the cluster centroid was selected as its representative. These 485 representative Korea10K samples were merged with 2,541 unrelated 1KGP individuals, yielding 517,041 high-quality, LD-pruned variants for ADMIXTURE analysis. Ten-fold cross-validation was used to determine the optimal K, defined as the model with the lowest cross-validation error (Fig. S2).

### Y-chromosome haplogroup classification and phylogenetic reconstruction

For 5,975 male Korean participants, Y-chromosome variants were lifted to GRCh37 using GATK LiftoverVcf v4.6.1.0 and analyzed with yhaplo v2.1.14.dev7 + gf8b300b (*47*) based on the ISOGG 2016 phylogeny. Of these, 104 failed to detect appropriate haplogroups being assigned to A major haplogroup, leaving 5,871 individuals with successful classification. Of 5,350 unrelated males, 60 failed. A total of 5,290 were used to calculate Y-haplogroup proportion in Korea.

Cross-population comparisons were performed using 1,244 male samples in the 1000 Genomes Project (1KGP) Phase 1 dataset.

For phylogenetic reconstruction, pairwise genetic distances were computed using VCF2Dis v1.54 (*48*) based on p-distance of VCF containing 5,871 haplogroup-classified males. To assess the robustness of the topology, we generated 100 bootstrap replicates, each constructed by randomly sampling 25% of SNPs. For each replicate, a Newick tree was inferred, and the resulting 100 trees were merged to obtain a final consensus tree. Consensus tree reconstruction and visualization were performed using IQ-TREE3 v3.0.1 (*49*).

### Mitochondrial DNA assembly and quality control

Filtered mitochondrial reads (chrM) were extracted from whole-genome BAM files using samtools v1.18 and converted to paired-end FASTQ format. Mitochondrial genomes were then assembled using NOVOPlasty v4.3 (50), with parameters optimized for mitochondrial assembly (genome range = 12,000–21,000 bp; k-mer = 39; maximum memory = 20 GB; insert size = 300 bp; read length = 151 bp; Illumina paired-end reads). Ambiguous nucleotides (Reduce ambiguous N’s) were reduced, and automatic insert size estimation was enabled.

A total of 10,239 samples were processed, from which 73 assemblies were excluded due to non-circularized sequences or multiple contigs. The final dataset comprised 10,166 high-quality circularized mitochondrial genomes. The rest of the mitochondrial genomes were generated making a consensus genome of the mitochondrial variants to the rCRS reference sequence using bcftools v1.21 (consensus) (51).

Annotation of 10,166 circularized assemblies to check their completeness was performed using MitoZ v3.5 (52) with the following parameters: annotate --species_name "Homo sapiens" --genetic_code 2 --clade "Chordata".

### Mitochondrial haplogroup classification and phylogenetic reconstruction

Mitochondrial haplogroups were assigned to 10,239 individuals via Haplogrep3 v3.2.1 (53) with the PhyloTree Build 17.2 database (phylotree- rcrs@17.2) .

All 10,166 high-quality circularized mitochondrial genomes were collected together with the revised Cambridge Reference Sequence (rCRS) and the Reconstructed Sapiens Reference Sequence (RSRS) for phylogenetic analysis. To ensure consistent alignment, each sequence was rotated to match the rCRS starting point at the first 30 bp of the CO2 gene, allowing up to four mismatches by using rotate v1.0 (54). Multiple sequence alignment was conducted using MAFFT v7.525 (55) with the --auto parameter.

Phylogenetic tree reconstruction was performed using IQ-TREE3 v3.0.1. An initial model search was conducted with ModelFinder (MFP), and the best-fitting substitution model identified in this step was subsequently applied to the final analysis. The maximum-likelihood tree was inferred under the TIM3+F+I+R6 model with 1,000 ultrafast bootstrap replicates (-B 1000) and 1,000 SH-aLRT tests (-alrt 1000). All analyses were run using 32 computational threads (-T 32).

Divergence time estimation was subsequently conducted in MEGA12 (56) using the RelTime framework. For this analysis, we imported the IQ-TREE Newick file and applied four previously reported calibration constraints corresponding to haplogroups A, B, C, and D, each defined by established time intervals (57). RelTime analysis was implemented under a maximum-likelihood statistical model.

To characterize rate variation across sites, we employed a General Time Reversible (GTR) nucleotide substitution model with gamma-distributed rate heterogeneity and a proportion of invariant sites (G+I). Rate variability was approximated using four discrete gamma categories. Sites with excessive missing data were filtered using partial deletion, retaining positions with at least 95% coverage. All RelTime computations were performed with 3 threads, enabling efficient processing of the large mitochondrial dataset.

### HLA Typing from Whole-Genome Sequencing Data

High-resolution human leukocyte antigen (HLA) genotyping was performed on 10,239 Korean whole-genome sequencing samples. Prior to typing, BAM files were normalized to ensure consistent alignment within the HLA region, particularly for samples generated with ALT or HLA contigs or non-standard alignment parameters, as advised by xHLA. Following the procedure implemented in get-reads-alt-unmap.sh, reads were extracted using samtools v1.13 (58) by selecting (i) unmapped reads or reads with unmapped mates, (ii) reads mapping to the canonical HLA locus on chromosome 6 (chr6:29,844,528–33,100,696), and (iii) reads mapping to all HLA-associated or ALT contigs (HLA- or chr6 alt-contigs).

Extracted reads were converted to FASTQ and re-aligned with BWA-MEM v0.7.17 using default parameters to the GRCh38 chromosome 6 primary assembly. The resulting BAM was sorted and indexed, and only the chr6:29.8-33.1 Mb region was retained for downstream analysis. HLA genotyping was conducted using xHLA (https://github.com/humanlongevity/HLA). HLA alleles were reported as classical two-digit resolution.

Only 9,000 unrelated individuals (therefore, 18,000 haplotypes and corresponding HLA-A pairs) were used to calculate the proportions of HLA-A alleles in Korean.

### Identification and Regional Aggregation of CG Context-Associated Variants (CGVs) for Population-specific profiling

Genomic variants altering local CG dinucleotide context were identified by comparing the flanking nucleotides of reference and alternate alleles in GRCh38 to generate a CG context index. Variants whose substitution altered this index were classified as CG context-associated.

To estimate the prevalence of CG context-associated variants (CGVs) across the Korean population, Population prevalence was assessed via allelic presence, scoring heterozygous or homozygous alternate genotypes as 1 and homozygous reference as 0. Variants present in ≥80% of individuals were defined as Korean-popular CGVs.

To quantify their impact on genomic CG composition, Korean-popular CGVs were mapped to 10-kb windows. Net gain or loss of CG sites per window was summed and normalized by the total CG count in GRCh38, producing a CG context change ratio. Windows with ≥10% net gain or loss were retained for downstream analysis.

### Comparing Allele Frequencies of CGVs Across Populations

To evaluate population-specific differences in CGVs, allele frequencies were compared between Koreans and individuals of European ancestry from the 1000 Genomes Project (1KGP). Variant allele counts were obtained using PLINK2 without prior biallelic filtering. Multiallelic variants were decomposed into biallelic representations and left-normalized using bcftools v1.21 for co-ordinate consistency with the Korean dataset.

For genomic windows in Koreans exhibiting ≥10% CG context alteration, we selected variants whose directional effects matched the overall trend in that region (i.e., variants increasing CG content in gain regions or reducing CG content in loss regions). Differences in allele counts between Korean and European populations were assessed using chi-squared tests. Resulting P-values were adjusted for multiple comparisons using the Benjamini–Hochberg false discovery rate (FDR) method.

### Genomic Annotation and Functional Enrichment of Hotspot CGVs

Genomic intervals exhibiting substantial CG context alteration were annotated using Annotatr (59) with the “hg38_cpgs”, “hg38_basicgenes”, and “hg38_enhancers_fantom” annotations to map proximal genes and regulatory features. Enrichment analyses for Gene Ontology (GO) terms and pathway associations were performed using ShinyGO (60) and GeDiPNet (61). Significance of enrichment was assessed using FDR correction, with terms considered significant at FDR < 0.05.

### Harmonization of Clinical Biomarkers Across KGP and NHANES Cohorts

Health check-up (HC) records were obtained from participants in the Korean Genome Project (KGP) and from publicly available datasets in the U.S. National Health and Nutrition Examination Survey (NHANES) III and IV. NHANES data were retrieved using the BioAge R toolkit (Kwon et al., 2021), which provides harmonized clinical and mortality records released by the U.S. Centers for Disease Control and Prevention.

Raw clinical data from KGP participants (N = 6,583) were harmonized with NHANES III (N = 18,825) and NHANES IV (N = 40,839) using a unified naming and unit-conversion system for clinical biomarkers. To ensure comparability across cohorts, biomarkers exhibiting substantial missingness were removed. To guarantee statistical power for genome-wide association analysis, only biomarkers representing at least 5,000 KGP participants were retained, except lymphocyte and monocyte proportions (deliberately included for possible adjustments for cell-type composition). A total of 12 clinical biomarkers were included in the mortality modeling: blood creatinine, waist circumference, body-mass index (BMI), blood triglyceride, chronological age, blood glucose, systolic blood pressure, diastolic blood pressure, high-density lipoprotein, total cholesterol level, lymphocyte proportions, and monocyte proportions.

Finally, only samples with complete phenotypic and consistent mortality status data were included in downstream analyses (NHANES III: N = 11,562, NHANES IV: N =16,173, and KGP: N = 4,498).

### Mortality Risk Prediction and Selection of Mortality-Associated Biomarkers

Candidate mortality-associated biomarkers were identified from NHANES III using penalized Cox proportional hazards regression implemented in the lifelines v0.30.0 (Python package), applying elastic-net regularization (α = 0.5). Each clinical variable was first examined by univariate Cox regression to assess the significance and direction of its hazard ratio. A multivariate penalized model was fitted to evaluate joint contributions of variables to mortality risk. Model performance was evaluated based on the concordance index (C-index). The resulting set of mortality-associated biomarkers, including creatinine, chronological age, glucose, and systolic blood pressure, was retained for downstream survival modeling.

### Multivariate Modeling of Phenotypic Age (PhenoAge)

Phenotypic Age (PhenoAge) is defined as the chronological age at which a reference individual exhibits the same predicted 10-year mortality hazard as that derived from a participant’s biomarker profile. Our model derives biological aging from clinical biomarkers only, not DNA methylation measures.

PhenoAge was calculated using a modified implementation of the Levine et al. (2018) algorithm. The BioAge R toolkit, specifically the “phenoage_nhanes” function, enabled the incorporation of the tailored biomarker panel determined from univariate mortality-risk modeling. A Gompertz proportional hazards model was fitted on NHANES III mortality data, linking the mortality-associated biomarkers to 10-year mortality risk.

The trained Gompertz model parameters were subsequently applied to the KGP dataset to generate PhenoAge estimates for Korean individuals. To align PhenoAge with chronological age and correct for regression bias, a linear calibration was performed.

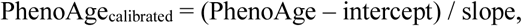

where the slope and intercept were obtained from linear regression of PhenoAge versus chronological age. This transformation ensured a mean PhenoAge Advance (PAA = PhenoAge – chronological age) centered around zero at the population level.

### Genome-wide Association Study on Phenotypic Age Advance (PAA)

From 9,000 unrelated Koreans, variants were retained after applying the following exclusion criteria: (i) minor allele frequency (MAF) < 1%, (ii) deviation from Hardy-Weinberg equilibrium (HWE) with *P* < 10^-6^, and (iii) missing genotype rate > 1%, leaving 6,996,863 variants.

Phenotypic Age Advance (PAA) exhibited a strong right-skewness and heavily-tailed distributional properties, violating linear regression assumptions. We applied a rank-based inverse normal transformation (INT) prior to association testing, yielding PAA-INT.

GWAS was performed on PAA-INT using PLINK2 (under a linear additive model) in 4,498 unrelated Koreans with matched genotype and clinical data. Covariates included chronological age, age^2^ (age squared), and the top 10 principal components (PC1-PC10) derived from genome-wide SNVs to adjust for the effect of time (linear and non-linear) and residual population stratification, respectively.

## Author contributions

Conceptualization: KA, SJ, YK, JB, GMC, NHP, HJ. Methodology: KA, SJ, YK, YC, CY, HJC, HL, YJK. Software: KA, SJ, YK, YC, CY, YJeon, JJeon. Formal Analysis: KA, SJ, YK, YC, CY. Investigation: KA, SJ, YK, YC, CY, JJeon, DS, HJC, HL, HR, AB, ESS. Resources: ESS, DB, SYLee, WK, MGK, AYH, SC, JTW, SYR, SiwooLee, HJBan, HJJin, YB, YMA, SJR, MJK, SYLee2, CMY, SHS, SJC, SGK, HTJ, BJH, YYChoi, JHCheong, SKKim, JHP, SAChoi, HYG, SYJ, JJung, WShin, SHLee, BK, WMyung, CKCheon, DUK, SSB, GNJ, HojuneLee, KSChae, CGKim, BHLee, JLee, KWKim, SL. Data Curation: ESS, DB, NHP, SiwooLee, HJBan, HJJin, YB, YMA, SJR, MJK, SYLee2, CMY, SHS, SJC, SGK, HTJ, BJH, YYChoi, JHCheong, SKKim, JHP, SAChoi, HYG, SYJ, JJung, WShin, SHLee, BK, WMyung, CKCheon, DUK, SSB, GNJ, HojuneLee, KSChae, CGKim, BHLee, JLee, KWKim, KA, SJ, YK, YC, CY, HJC, HL, YJeon, JJeon, SJoe, JOY, JHKim. Visualization: KA, SJ, YK, YC, HJC, HL. Writing - Original Draft: KA, SJ, YK, YC. Writing - Review & Editing: CY, HJC, HL, YJK, DS, HR, AB, SP, JC, SL, BCK, JK, DB, YCho, KC, SYLee, WK, MGK, AYH, SC, JJung, WShin, SHLee, BK, WMyung, CKCheon, DUK, SSB, GNJ, HJL, KSChae, CGKim, BHLee, JLee, KWKim. Supervision: NHP, HJ, GMC, JB. Project Administration: NHP, HJ, GMC, JB, YJeon, JJeon, SJoe, JOY. Funding Acquisition: JB, SJ HJ, HR, YJeon, JJeon, SJoe, JOY.

## Competing interests

SJ is CEO and HR is an employee of AgingLab Inc. YCho is an employee of CG Invites Co., LTD and Invites Genomics Co., LTD. B.L. is CEO and JL is an employee of nSAGE. GC is co-founder of Nebula Genomics and Glottatech.com. All other authors declare no competing interests.

## Acknowledgments

We appreciate all participants and Ulsan citizens who supported this project. We thank former members of Korean Genomics Center (KOGIC), namely Hansol Choi, Whan-Hyuk Choi, Muhyeok Kim, Chanyoung Lee, Jeongwoo Seo, Hyeonjae Lee, Seolbin An, Hyeonju Choi, Hyeonsu Cho, Jiyu Lee, Hwan Yu, Gwon Yul Jo, Changjae Kim, Yeonkyung Kim, Younghui Kang, Yeong Ju Woo, Jungae Shim, Nayeong Kim, Shinseob Yoon, Suji Hong, Ju Yeon Park, Boram Park, Sangryoul Han, Yeshin Park, Hak-Min Kim, Suan Cho, and Seung Gu Park who contributed to this project. We thank Moo-Jae Cho and Moo-Young Jung of Ulsan National Institute of Science and Technology (UNIST). We thank Du-Gyeom Kim, Cheol-Ho Song, Ki-Hyun Kim, Jung-Ik Kim, Soo-Jung Shin, Byung-Ryel Yoo, Jeong-Sik Yang of Ulsan City for their support on this project. This work was supported by Biodatafarm computing infrastructure funded by the Ulsan metropolitan city government. We thank Seungwoo Jeon, Changwoo Park, Doyung Oh for supporting the use of Biodatafarm. We thank the Korea Institute of Science and Technology Information (KISTI) for providing us with the Korea Research Environment Open NETwork (KREONET). We thank our collaborators in National Center for Standard Reference Data (NCSRD) of the Korea Research Institute of Standards and Science (KRISS).

The biospecimens and data used in this study were provided by: the Biobanks of Ulsan University Hospital (60SA2016001-002, 60SA2016001-003, 60SA2016001-005, 60SA2016001-010-5, 60SA2017002-001, 60SA2017002-004, and 60SA2017002-006-4); Ulsan Medical Center; the Biobanks of Gyeongsang National University Hospital (KHUH 2018-11-043); the Biobanks of Kyung Hee University Hospital (2018-4, 2019-4, 2019-6, 2021-2); Chungbuk National University Hospital (18-27, 20-4, 21-3), a member of the National Biobank of Korea, which is supported by the Ministry of Health, Welfare and Family Affairs; the Biobank of Pusan National University Hospital a member of the Korea Biobank Network (19092301-08-01); the support of Borah Kim, MD, Ph, Assistant Professor, Department of Psychiatry, CHA Bundang Medical Center, CHA University (CHAMC 2018-11-004); the Biobank of Severance Hospital, Seoul, Korea. All samples derived from the National Biobank of Korea were obtained with informed consent under institutional review board-approved protocols

This work was supported by the U-K BRAND Research Fund (1.200108.01) of UNIST (Ulsan National Institute of Science & Technology), the Research Project Funded by Ulsan City Research Fund (1.200047.01) of UNIST, the Promotion of Innovative Businesses for Regulation-Free Special Zones funded by the Ministry of SMEs and Startups (MSS, Korea) (1425157301 and 1425156792), the Establishment of Demonstration Infrastructure for Regulation-Free Special Zones funded by the Ministry of SMEs and Startups (MSS, Korea) (1425157253), and the Technology Innovation Program (20016225, Development and Dissemination on National Standard Reference Data) funded by the Ministry of Trade, Industry & Energy (MOTIE, Korea). This work was also supported by funding from the Korea Planning & Evaluation Institute of Industrial Technology with support from the Ministry of Trade, Industry and Energy in 2024 [RS-2024-00435468, Development and Dissemination of National Standard Technology.

We also acknowledge the use of ChatGPT (OpenAI) for assistance with language editing, polishing, and improving the clarity of the manuscript.

## Notes

### Summary of Updates

Authors and Affiliations updated; Figure 5 and According Results added; Supplementary Figure 4 revised; Main text of the manuscript revised

