## Supplementary Materials for "10,239 whole genomes with multiomic and clinical health information as the Korean Multiomics Reference dataset"

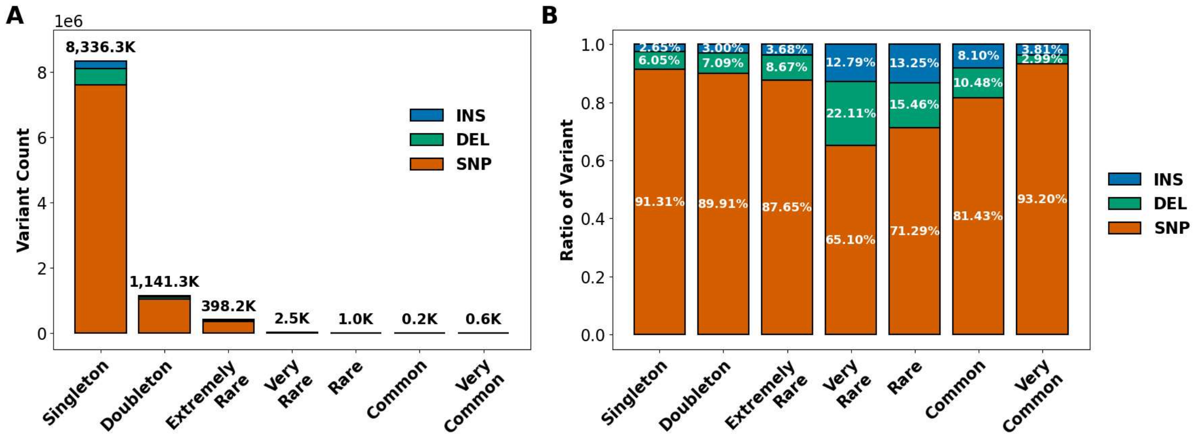


**Fig. S1. Characteristics of novel variants across allele frequency categories.** Bar plots showing **(A)** count and **(B)** proportion of novel variants (y-axis) across allele frequency bins (x-axis) stratified by variant type: SNP (orange), deletion (DEL; green), and insertion (INS; blue). Variant counts are shown as stacked bars with absolute values annotated above, while proportions are displayed as relative contributions within each frequency bin.

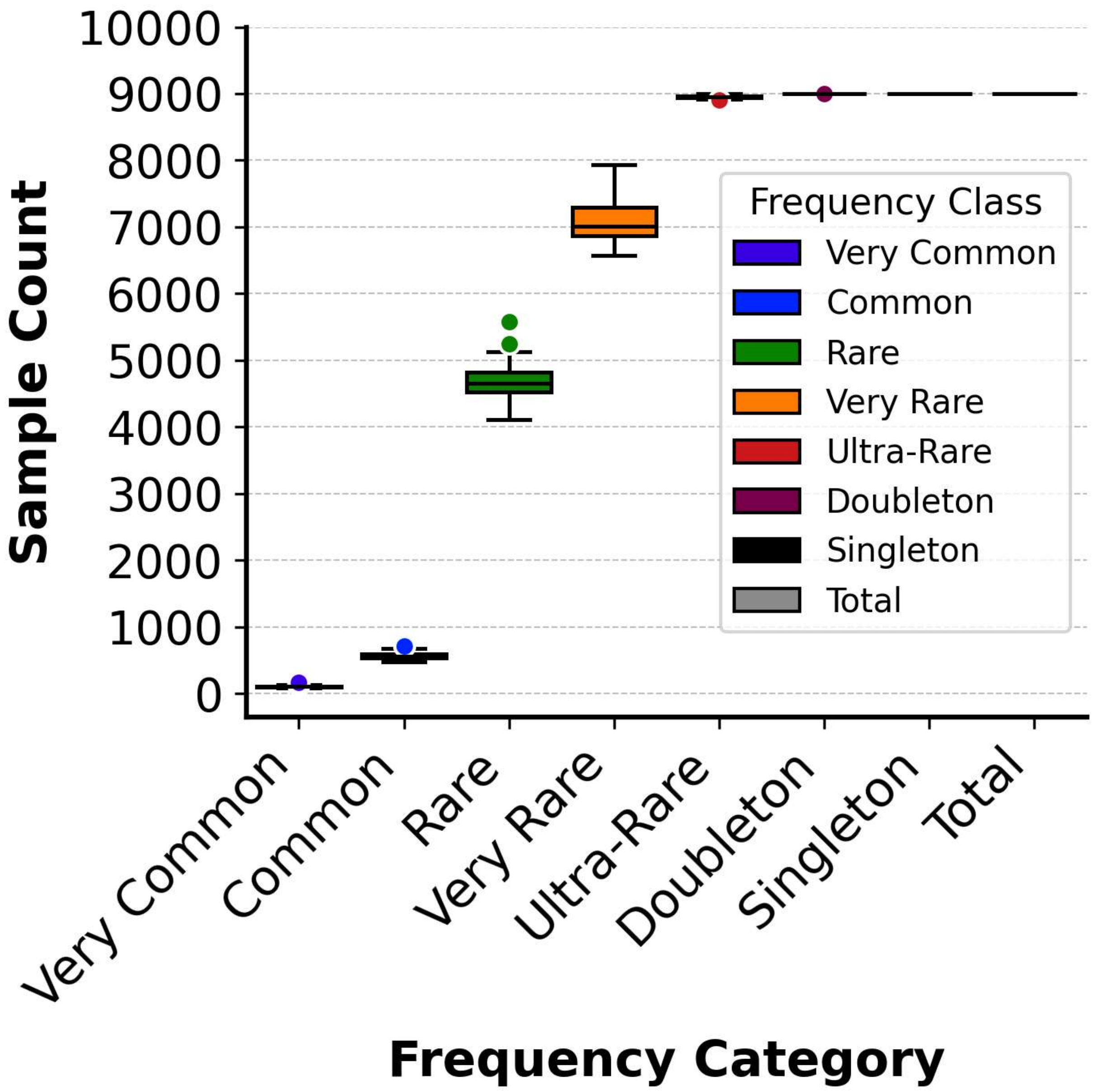


**Fig. S2. Saturation analysis across allele frequency categories in 9,000 unrelated Koreans.** Boxplots depict the distribution of sample counts for variant discovery (y-axis) stratified by minor allele frequency (MAF) category (x-axis). Each box represents the interquartile range (IQR, $25^{th}$-$75^{th}$ percentile) with the median indicated by a horizontal line; whiskers extend to 1.5×IQR. Individual variants beyond the whiskers are shown as colored dots corresponding to frequency class. Sample counts reaching 9,000 samples, as exemplified by singleton and doubleton, indicates failure of the saturation in the cohort.

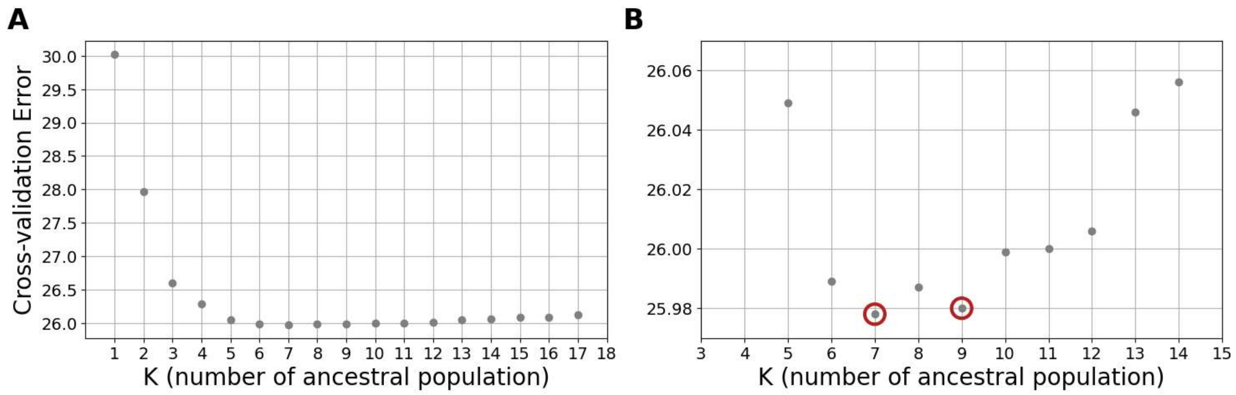


**Fig. S3. Selection of Optimal ADMIXTURE K-value(s) with minimal cross-validation errors. (A)** Scatter plot showing the cross-validation (CV) error (y-axis) and K (number of hypothetical ancestral populations; x-axis). The CV error decreases sharply, reaching a plateau around K=7, suggesting this value captures the major structure in the dataset. **(B)** Magnified view of the cross-validation error between K = 3 and K = 15. The plot highlights two distinct local minima at K = 7 and K = 9 (indicated by red open circles).

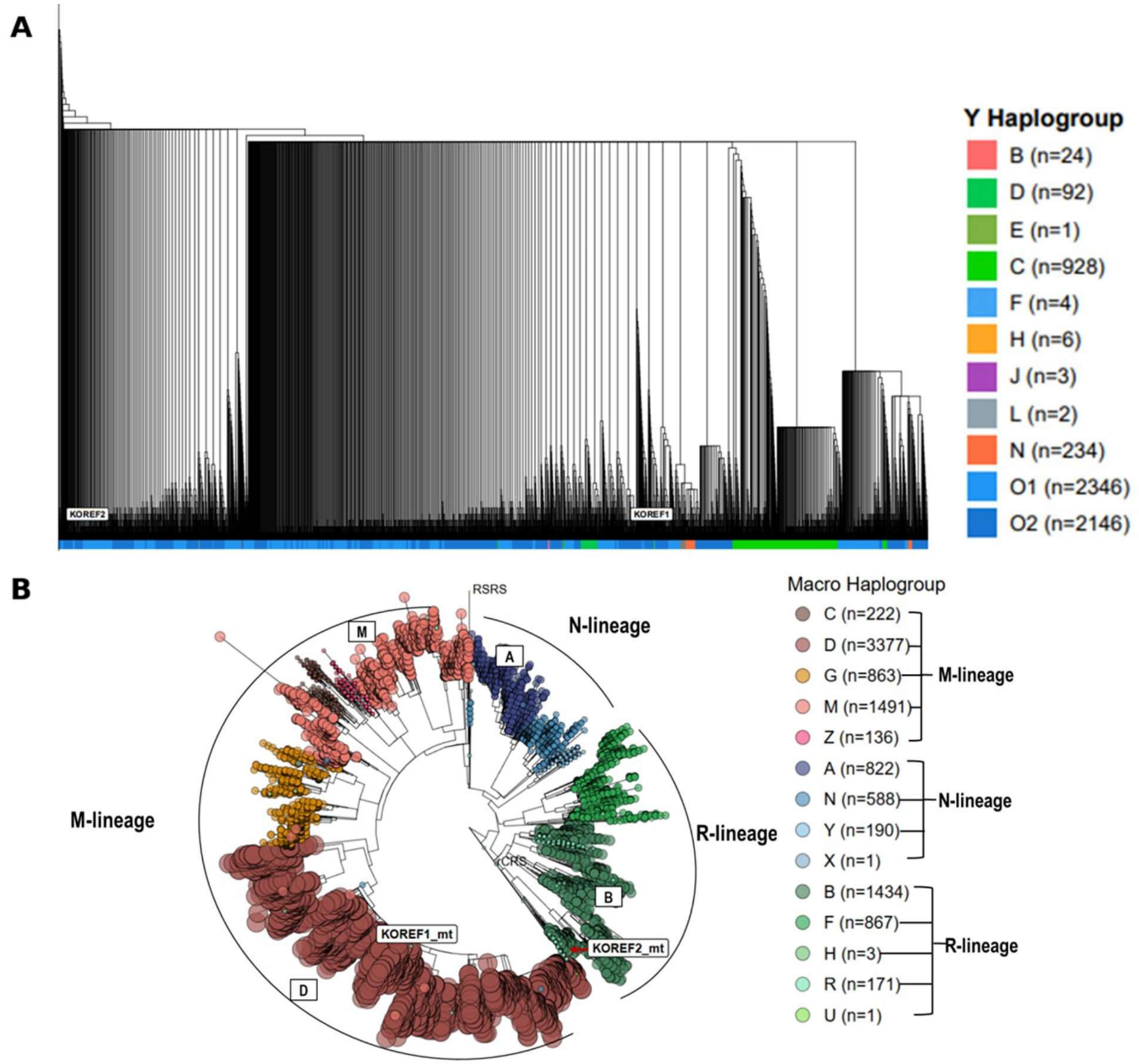


**Fig. S4. Phylogenetic reconstruction of Korean uniparental lineages in Korea10K. (A)** A Y-chromosomal lineage tree based on p-distance showing 5,871 Y haplogroup-classified Korean males. **(B)** A mitochondrial lineage tree based on maximum likelihood portraying 10,166 MT haplogroup-assigned Korean individuals. Reference sequences, including Korean (KOREF1 and KOREF2), European (rCRS), and hypothetical human ancestor (RSRS) are annotated within the trees.

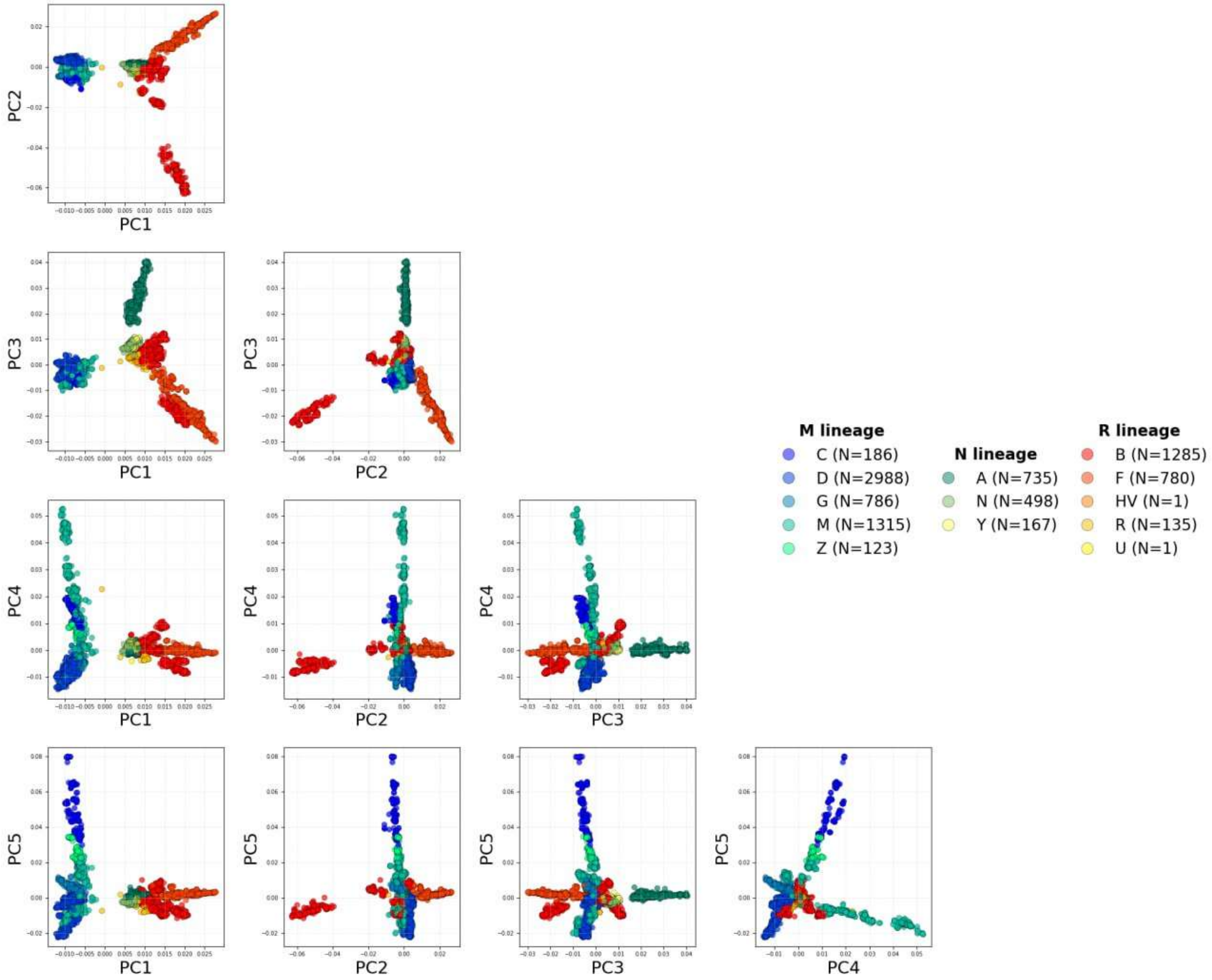


**Fig. S5. Combinatorial PC-biplots of mitochondrial variants in Korea10K.** Each biplot of PCs depicts the distribution of mitochondrial variants along principal component axes (from PC1 to PC5 in a combinatorial manner), highlighting sample clustering according to haplogroup assignments. Colors indicate ancestral lineages (macrohaplogroups) corresponding to each haplogroup.

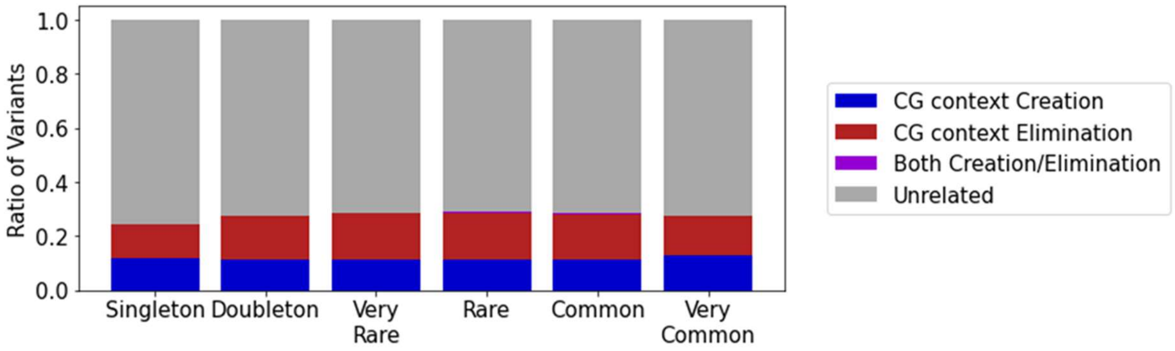


**Fig. S6. Bar plot depicting the characteristics of CGVs across allele frequency categories.** Proportion of CGVs (y-axis) across allele frequency bins (x-axis) categorized by CG context impact: creation (blue), elimination (red), both creation/elimination (purple), and unrelated (gray).

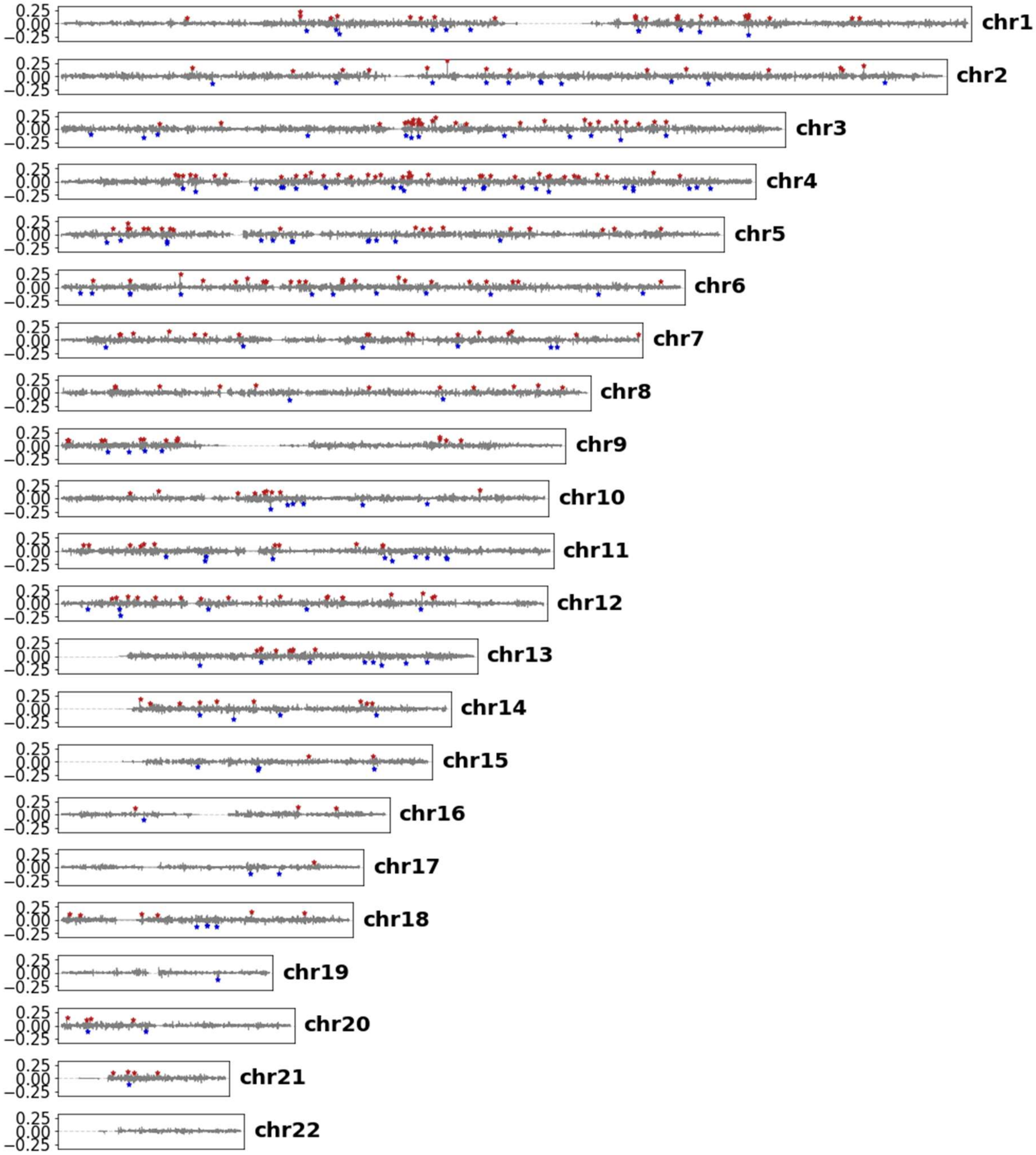


**Fig. S7. Ratio difference of CG context by chromosome region.** Visualization of the proportion of CG context changes (y-axis) across 10,000 bp windows for each chromosome in GRCh38 (x-axis). The y-axis represents the relative difference (ratio) of CG context compared to GRCh38. Regions showing more than a 10% change are marked with stars.

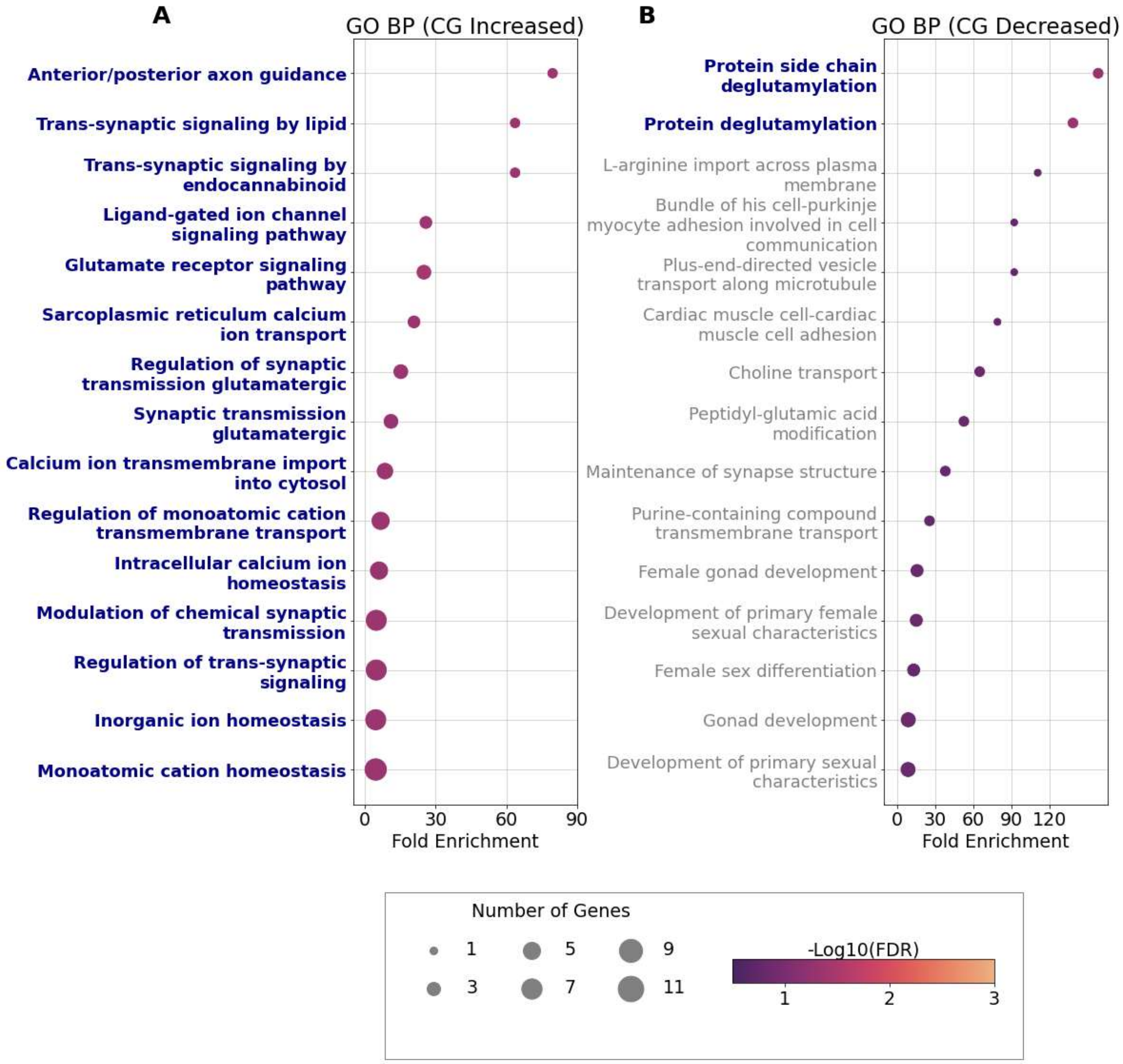


**Fig. S8. Gene set enrichment analysis of regions with more than 10% relative change in CG context (GO Biological Process). (A)** Regions with a relative increase over 10%; **(B)** Regions with a relative decrease over 10%; Only the top 15 enriched terms are shown. The x-axis represents fold enrichment, while the colors indicate significance (negative of log10*FDR*, lighter = more significant). The y-axis lists GO terms in GO Biological Process. The names of terms with a significant false discovery rate (*FDR* < 0.05) are shown in blue.

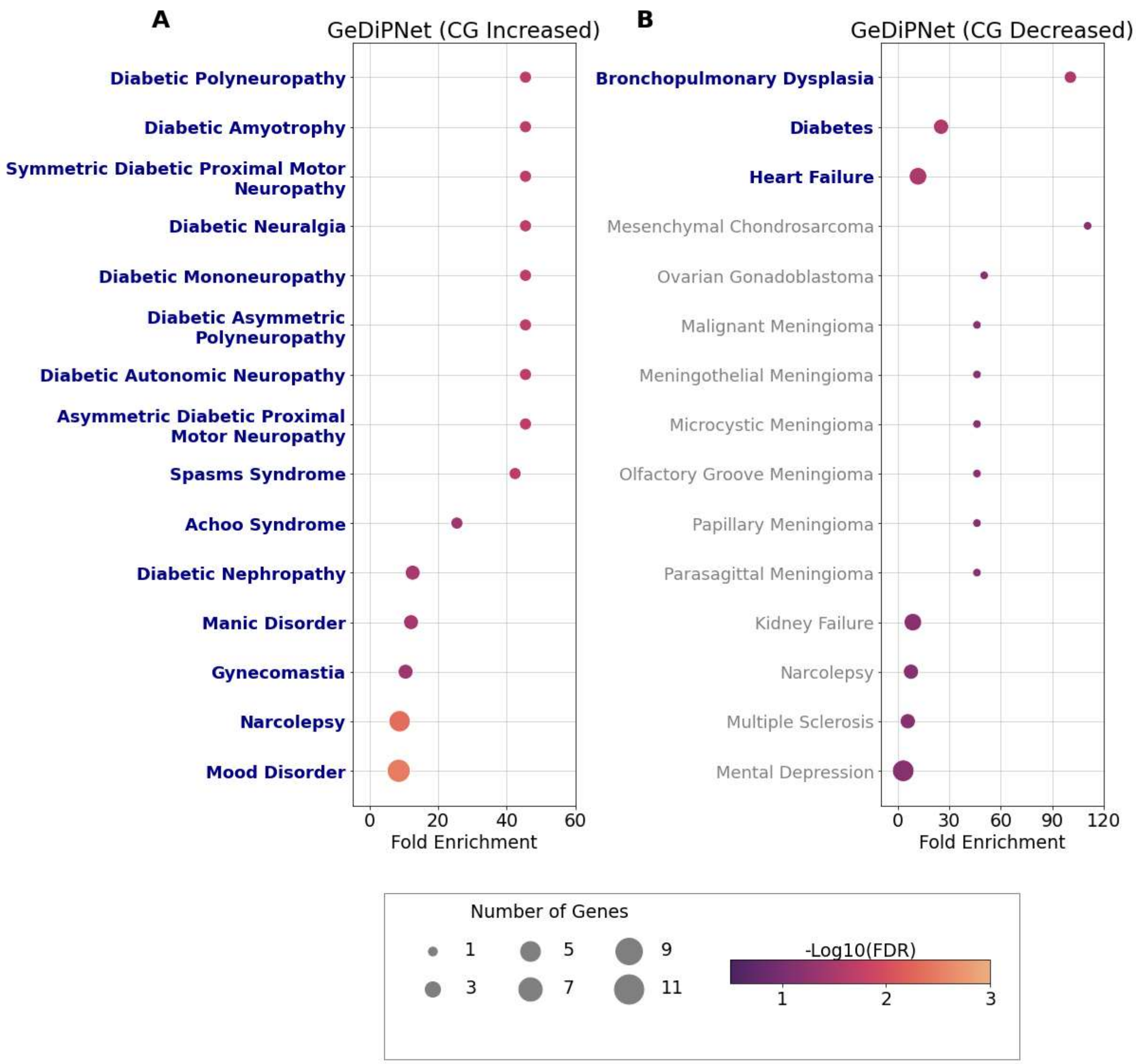


**Fig. S9. Gene set enrichment analysis of regions with more than 10% relative change in CG context (GeDiPNet). (A)** Regions with a relative increase over 10%; **(B)** Regions with a relative decrease over 10%; Only the top 15 enriched terms are shown. The names of terms with a significant false discovery rate (*FDR* < 0.05) are shown in blue. The x-axis represents fold enrichment, while the colors indicate significance (negative of log10*FDR*, lighter = more significant). The y-axis lists GenDiPNet terms in GenDiPNet Biological Process. The names of terms with a significant false discovery rate (*FDR* < 0.05) are shown in blue.

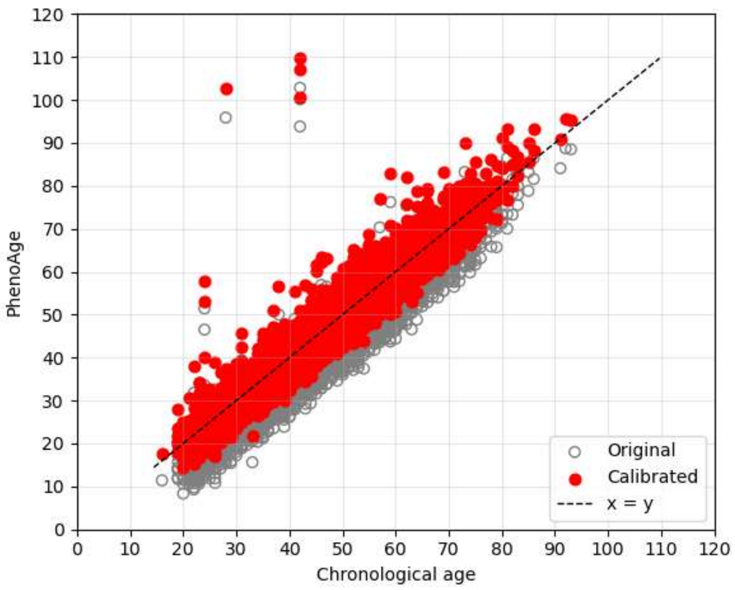


**Fig. S10. Calibration of PhenoAge to chronological age.** Predicted phenotypic age (PhenoAge) is plotted against chronological age for all individuals, with each point representing a single participant. Grey circles show the original PhenoAge estimates, and red circles show the calibrated PhenoAge values after linear recalibration to the chronological age distribution. The dashed line indicates the identity line (x = y), along which regression line for calibration.

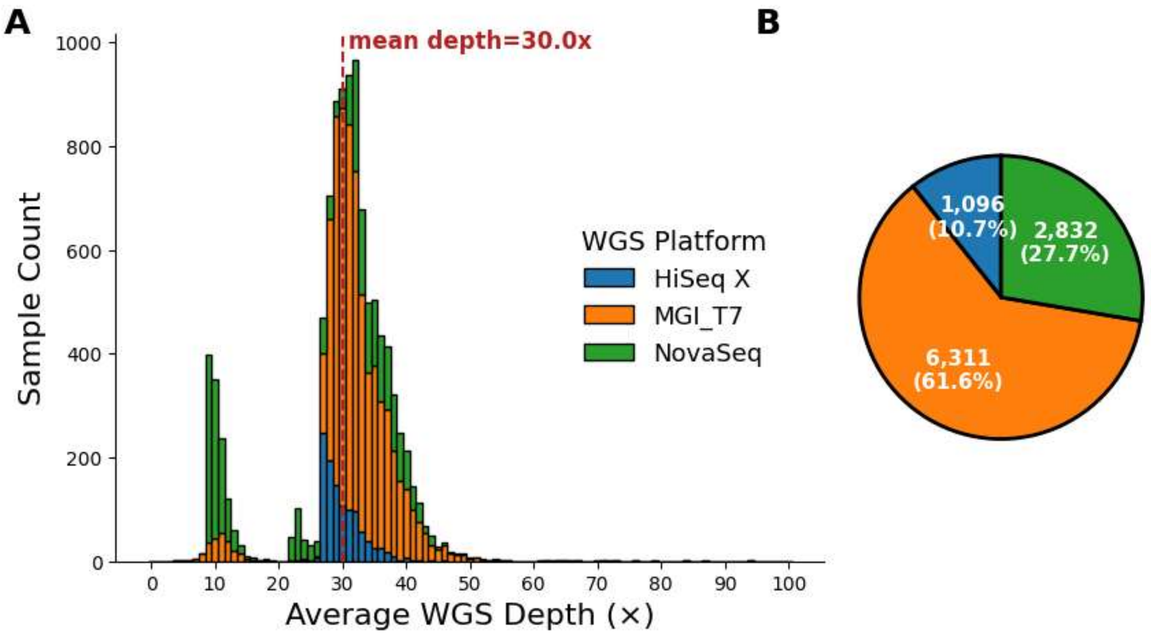


**Fig. S11. Distributions of sequencing depths and platforms across 10,239 whole genomes.** **(A)** Stacked bar plot showing the distribution of average sequencing depth per sample (x-axis, as ×) across 10,239 Korean genomes (y-axis). Samples are grouped by rounded average depth (0–100×) on the x-axis, with colors indicating WGS platform: HiSeq X (blue), MGI T7 (orange), and NovaSeq (green). The vertical red dashed line and annotation indicate the mean depth (30.0×). The y-axis denotes the number of samples. **(B)** Pie chart showing the proportion of samples sequenced on each platform, with absolute counts and percentages indicated on the wedges.

**Table S1. Summary statistics of saturation analysis for full variant discovery stratified by allele frequency.** The table shows the number of genomes required to reach saturation in variant detection across different frequency categories. For each category, the minimum (Min), median (Median), maximum (Max), and mean (Mean) number of genomes at saturation are reported, along with the 95% confidence interval (95% CI [lower, upper]) of the mean. Red highlights statistics reaching the maximum number (9,000 samples) without full variant discovery, which indicates failure to saturate.

**Table S2. Summary statistics of imputation performance stratified by allele frequency.** The table shows percentage improvement in imputation quality ($R^2$) was calculated for each allele frequency (AF) bin (%) by comparing imputation panels of Korea10K to Korea1K (10K/1K) and Korea10K to Korea4K (10K/4K).

**Table S3. Top five hotspot CGVs in Korea10K.** The table lists the genomic regions exhibiting the most pronounced changes in CG dinucleotide context relative to the GRCh38 reference genome. Columns indicate the chromosome, start and end coordinates of the 100 kb window, percentage change in CG content (% CG Difference), absolute number of CG changes (# CG Difference), total number of CG dinucleotides in the reference (# CG from GRCh38), and gene annotation or relevant genomic feature. Positive values indicate a gain in CG sites, whereas negative values indicate a loss.

**Table S4. Overview of Health Check-up (HC) measured in Korea10K.** The table summarizes the variables collected from participants across multiple domains including anthropometry, physiological and functional measures, hematology, and urinalysis. Normal reference ranges reflect commonly accepted clinical standards and are stratified by sex when appropriate.

**Table S5. Overview of Lifestyle Questionnaire (LQ) in Korea10K.** The table lists self-reported lifestyle variables, captured through structured questionnaires, that have been answered by survey respondents in Korea10K cohort.

**Table S6. Summary statistics of Whole-Genome Sequencing (WGS) data in Korea10K.** The table summarizes key sequencing metrics for Korea10K genomes. Columns include the sample identifier (SampleID), total sequencing output (Production_Amount, Gbp), calculated mean coverage depth (Average_Depth, ×), integer rounded coverage depth (Rounded_Depth, ×), and sequencing platform used (WGS Platform).
